# Ensemble-Based Modeling of the SARS-CoV-2 Omicron BA.1 and BA.2 Spike Trimers and Systematic Characterization of Cryptic Binding Pockets in Distinct Functional States : Emergence of Conformation-Sensitive and Variant-Specific Allosteric Binding Sites

**DOI:** 10.1101/2023.08.21.554193

**Authors:** Gennady Verkhivker, Mohammed Alshahrani, Grace Gupta

## Abstract

A significant body of experimental structures of the SARS-CoV-2 spike trimers for the BA.1 and BA.2 variants revealed a considerable plasticity of the spike protein and emergence of druggable cryptic pockets. Understanding of the interplay of conformational dynamics changes induced by Omicron variants and identification of cryptic dynamic binding pockets in the S protein are of paramount importance as exploring broad-spectrum antiviral agents to combat the emerging variants is imperative. In the current study we explore conformational landscapes and characterize the universe of cryptic binding pockets in multiple open and closed functional spike states of the Omicron BA.1 and BA.2 variants. By using a combination of atomistic simulations, dynamics network analysis, and allostery-guided network screening of cryptic pockets in the conformational ensembles of BA.1 and BA.2 spike conformations, we identified all experimentally known allosteric sites and discovered significant variant-specific differences in the distribution of cryptic binding sites in the BA.1 and BA.2 trimers. This study provided in-depth structural analysis of the predicted allosteric site in the context of all available experimental information, revealing a critical role and effect of conformational plasticity on the distribution and function of allosteric binding sites. The results detailed how mutational and conformational changes in the BA.1 and BA.2 pike trimers can modulate the functional role of druggable allosteric pockets across different functional regions of the spike protein. The results of this study are particularly significant for understanding the universe of cryptic bindings sites and variant-specific preferences for druggable pockets. Exploring predicted druggable sites can present a new and previously underappreciated opportunity for therapeutic intervention of Omicron variants through conformation-selective and variant-specific targeting of functional sites involved in allosteric changes.

## 1. Introduction

The unprecedented body of structural and biochemical studies have explored mechanisms of SARS-CoV-2 infection showing the key role of the SARS-CoV-2 viral spike (S) glycoprotein undergoing stochastic movements between distinct functional forms [1–9]. The complex architecture of the S protein includes an amino (N)-terminal S1 subunit experiencing functional motions and structurally rigid carboxyl (C)-terminal S2 subunit. Conformational transformations of the SARS-CoV-2 S protein between the closed and open S states are exemplified by coordinated global movements of the S1 subunit consisting of an N-terminal domain (NTD), the receptor-binding domain (RBD), and two structurally conserved subdomains SD1 and SD2 which together determine the structural and dynamic response of the S protein to binding partners and the host cell receptor ACE2 [10–15]. A number of biophysical studies provided in-depth characterization of the thermodynamic and kinetic aspects of the SARS-CoV-2 S functional trimer, showing the complex interplay between subdomain movements and long-range interactions that couple S1 and S2 subunits to modulate the RBD equilibrium and population-shifts between the RBD open (up) and closed (down) conformations regulating the exposure of the S protein to binding partners and strength of S-ACE2 binding [16–18]. The rapidly growing number of cryo-EM and X-ray structures of the SARS-CoV-2 S variants of concern (VOC’s) in various functional states and complexes with antibodies revealed a remarkable versatility of molecular mechanisms and diversity of binding epitopes that underlie binding affinity of S proteins with different classes of antibodies [19–28]. The population distribution of the closed and open states of these S trimers varies among different studies, due to subtle differences in the chemical condition used by different research groups. The cryo-EM structures of the S Omicron BA.1 trimer in the open and closed forms revealed that the dominantly populated conformation is the closed state with all the RBDs buried, leading to ‘conformational masking’ that may prevent antibody binding and neutralization at sites of receptor binding [24]. Another large scale cryo-EM study of the S Omicron BA.1 variants in different functional states showed tight packing of the RBDs in the Omicron 3-RBD-down structures that limits RBD motion, causing a prominent single protomer NTD-to-RBD (N2R) linker rearranged conformation [27]. The cryo-EM structures of the S Omicron BA.1 trimer showed that the Omicron sites N856K, N969K, and T547K can promote favorable interactions with D658, Q755, and S982 from neighboring subunits, resulting in the increased number of the inter-protomer contacts which can confer the enhanced stability [28]. Other studies evaluated the thermostability of the S-D614G, S-BA.1, and S-BA.2 protein ectodomains as well as the stability of their corresponding monomeric RBD constructs in the differential scanning fluorimetry (DSF) assays showing the reduced stability of the BA.1 RBD, while BA.2 RBD appeared to be more stable than BA.1 but less stable than the Wu-Hu-1 [29–32]. According to these studies, the remodeling of the interfacial RBD loops and corresponding enhanced interprotomer RBD-RBD packing in the 3-RBD-down form of the S-BA.2 trimer relative to the S-BA.1 can lead to further stabilization of the S-BA.2 protein [31,32]. Several structural investigations of the Omicron BA.1 variant presented evidence of the stabilization of the 1RBD-up open S conformation [33–35]. The reported 3.1 Å-resolution cryo-EM structure of the S Omicron protein ectodomain [34] and another 3.0 Å cryo-EM structure of the S Omicron protein ectodomain [35] showed that in contrast to the original strain of SARS-CoV-2 with a mixture of open and closed conformations, the S Omicron BA.1 proteins may adopt predominantly an open 1 RBD-up position predisposed for receptor binding. These studies indicated that S-BA.2 protein has a different population distribution of open and closed forms compared to the BA.1 protein. The population of spike proteins with one RBD in the up conformation is also higher for Omicron than for other variants, such as Delta, Kappa, and Beta variants [34,35]. Because more RBDs in the up conformation make the corresponding structure of the S protein more open for binding ACE2, the observed greater variability of the S-BA.2 trimers in the unbound and ACE2-bound forms may cause higher transmission and infection rates of the BA.2 Omicron sublineage compared to BA.1.The Omicron BA.2 subvariants of SARS-CoV-2 have been associated with the increased transmissibility and vaccine evasion capacity [45,46]. Structural and biophysical studies of the RBD-ACE2 complexes for the BA.1.1, BA.2, and BA.3 variants revealed that the binding affinity of the Omicron BA.2 with ACE2 is stronger than the affinities of the BA.3 and BA.1 subvariants [45]. Thermal shift assays showed that that RBD in Omicron BA.2 is more stable than that from BA.1, but the observed lower temperature for dissociation of the S-BA.2 trimer indicated that it is more dynamic than the S-BA.1 trimer, which may arise due to the inter-protomer salt bridges K856-D571 in the BA.1 and absence of these stabilizing interactions in the BA.2 trimer [46]. The structure of the S-BA.2 trimer complex with hACE2 revealed a more extensive interaction network in the RBD-hACE2 interface and combined with the higher stability of the BA.2 RBD compared to the BA.1 RBD might also contribute to the higher binding affinity of the BA.2 RBD to hACE2 [46].The recently reported structural and functional analysis of the S-BA.2 protein produced three distinct states representing the closed 3-RBD-down conformation, a 1-RBD-up conformation and an RBD-intermediate conformation, all of which were also found for the S-G614 trimer [47]. This study also highlighted reconfiguration of the NTD segment and the surface-exposed 143–154, 173–187, 210–217 and 245–260 loops several of which form important parts of the neutralizing epitopes in the NTD [47]. The reported cryo-EM structures of the S trimers for BA.1, BA.2, BA.3, and BA.4/BA.5 subvariants of Omicron showed that S-BA.1 trimer is stabilized in an open conformation while S-BA.2 exhibits two conformational states corresponding to a closed and an open form with one RBD in the up position [48]. The thermal stability assays demonstrated that S-BA.2 trimer was the least stable among BA.1, BA.2, BA.3 and BA.4/BA.5 variants revealing a less compact inter-protomer arrangements while BA.1, BA.3 and BA.4/BA.5 spike exhibited tighter inter-subunit organization with more buried areas between S2 subunits [48]. While numerous cryo-EM structures of S trimers of VOCs provided accurate high-resolution structural information and described the extent of opening of various variants, structures alone cannot uncover the intrinsic dynamics and conformational heterogeneity underlying S functions and binding. The hidden dynamic nature of the S trimers was elucidated using hydrogen–deuterium exchange monitored by mass spectrometry (HDX-MS) [49] uncovering an alternative open trimer conformation that interconverts slowly with the canonical prefusion structures and can dynamically expose novel epitopes in the conserved region of the S2 trimer interface, providing new epitopes in a highly conserved region of the protein. Another HDX-MS study we identified changes in the S dynamics for VOCs, revealing that Omicron mutations may preferentially induce closed conformations and that the NTD acts as a hotspot of conformational divergence driving immune evasion [50]. It was found that the divergence hotspot overlaps with the NTD antigenic supersite [51] and that the increased flexibility acquired by early VOC’s and Omicron variants promotes immune escape of NTD-targeting antibodies [52]. In particular, BA.2 displayed strong resistance to the vast majority of existing neutralizing monoclonal antibodies. A systematic comparison of efficacy for 50 human monoclonal antibodies covering the seven identified epitope classes of the SARS-CoV-2 RBD, against BA.1, BA.2, and BA.3 variants confirmed the enhanced immune evasion profile of the BA.2 variant which may arise from conformational variability of the S trimer[53,54]. These functional studies suggested that besides mutation-induced interaction changes, the variant-specific changes in the dynamic equilibrium of the S Omicron protein is an important factor affecting immune evasion profile. The recent studies showed that the conformation of the S protein determines plasma neutralizing activity elicited by human vaccines, mostly due to binding to the S1 subunit which comprises antigenic sites in the RBD and NTD recognized by most neutralizing antibodies [55]. It may be noted that while we consider in this study the conformational variability and intrinsic of the S trimer prefusion states, cryo–electron tomography experiments unveiled a far more complex landscape of the S protein refolding and captured partially folded intermediate spike states on the pathway to membrane fusion [56].The significant body of experimental structures of S trimers for BA.1 and BA.2 variants revealed a considerable plasticity of the closed and open forms and emergence of dynamic intermediate states. In addition, biophysical studies produced diverse and often conflicting data on the thermal stability of the S trimers for these variants, where most recent structures indicated that the BA.2 variant had a lower stability than the BA.1 and BA.2.75 variants. There is no consensus on how Omicron variants can affect thermodynamic preferences for the open or closed states, thus highlighting the elusive nature of dynamic spike equilibrium and allocation of binding sites that can be modulated by various mutations. A highly conserved cryptic epitope in the RBD was discovered in one of the early structural studies [57]. The cryo-EM structure of the S protein with linoleic acid (LA) complex revealed allosteric binding pocket in the S-RBD (often referred as LA pocket) that is modulated through opening of a gating helix and allosterically induces reduced levels of S binding in the presence of LA [58]. A highly conserved cryptic epitope at the S trimeric interface enables broad antibody recognition of Omicron variants suggesting a possible mechanism for antibody neutralization via inducing S trimer disassembly [59]. A cryptic binding pocket in the NTD was revealed in structural studies showing that polysorbate (PS) detergent molecule can occupy this site when detergent was present in the formulation of the immunogen [60]. Cryo-EM and X-ray crystallography studies discovered that this cryptic site on the NTD binds the tetrapyrrole products of heme metabolism, biliverdin and bilirubin, with nanomolar affinity blocking antibody access to their NTD epitopes on the S protein [61]. The cryo-EM structure of a vaccine-induced antibody discovered a novel neutralizing epitope on the NTD that is defined by a cryptic hydrophobic NTD cavity that binds a heme metabolite biliverdin [62]. Moreover, due to a generally conserved and yet plastic nature of the epitope, the antibody maintains binding to VOC’s including Omicron BA.1 variant [62]. Other latest studies highlighted conformational plasticity of the NTD regions where mutations/deletions not only change the architecture, but also alter the surface properties, leading to remodeling of the binding pockets and major antigenic changes in the NTD supersite and loss of antibody binding [63]. These studies support the recent data suggesting that the NTD of the S protein can serve as an adaptable antigenic surface capable of unlocking and redistributing cryptic binding pockets which may enable the virus to divert the host immune responses away from the RBD regions [50].Computer simulations provided important atomistic and mechanistic advances into understanding the dynamics and function of the SARS-CoV-2 S proteins. [64–70]. Conformational dynamics studies using cryo-EM showed that the closed ground S state may be in equilibrium with 1 RBD-up form that is trapped by ACE2 binding leading to shifting of the conformational landscape of S trimer and enhanced conformational plasticity in S1 subunits [67]. Large scale adaptive sampling simulations of the viral proteome captured conformational heterogeneity of the S protein and predicted the existence of multiple cryptic epitopes and hidden allosteric pockets [68]. The replica-exchange molecular dynamics (MD) simulations examined conformational landscapes of the full-length S protein trimers, discovering the transition pathways via inter-domain interactions, hidden functional intermediates along open-closed transition pathways and previously unknown cryptic pockets that were consistent with FRET experiments [69,70]. The cryo-EM MetaInference (EMMI) method modeled conformational ensembles by combining simulations with cryo-EM data, revealing the intermediate states in the opening pathway of the S protein and discovering potentially druggable novel cryptic sites near the RBD recognition site [71]. Our recent studies combined multiscale simulations with coevolutionary analysis and network-based modeling to reveal that the S protein can function as a functionally adaptable allosterically regulated machine that exploits plasticity of allosteric centers to fine-tune response to antibody binding [72–76].

Computer simulation-based mapping of the full-length models of the S protein in the presence of benzene probes reproduced experimentally discovered allosteric sites and identified a spectrum of novel cryptic and potentially druggable pockets [77]. MD simulations of the unbound or ACE2-bound RBD conformations combined with pocket analysis and druggability prediction identified several promising druggable sites, including one located between the RBD monomers [78]. Recent functional studies identified that the metabolite of fenofibrate (fenofibric acid or FA) destabilized the RBD protein and significantly reduced infection rates in vitro by inhibiting the ACE2 binding [79]. MD simulations and energetic analysis suggested that the FA induces a conformational change in the RBD by stabilizing a potential cryptic binding site on the RBD [80]. The reversed allosteric communication approach based on the premise that allosteric signaling in proteins is bidirectional and can propagate from an allosteric to orthosteric site and vice versa, has been used for characterization of binding shifts in the protein ensembles and identification of cryptic allosteric sites [81–83]. A network-based adaptation of the reversed allosteric communication approach identified allosteric hotspots and RBD binding pockets in the Omicron variant RBD-ACE2 complexes [84]. Integration of computational and experimental studies enabled discovery and validation of the cryptic allosteric site located between subdomains of the S protein, with several compounds targeting this site showing characteristic binding and anti-virus activities [85,86].A large number of binding pocket predictors is available [87–91] including the growing number of methods based on exploring the dynamic protein ensembles [92–95] as well as rapidly growing tools empowered by artificial intelligence (AI) and machine learning (ML) approaches [96–111]. There has been the increasing interest in identifying druggable sites in the S2 subunit as most recently developed broad-spectrum fusion inhibitors and candidate vaccines can target the conserved elements in the S2 subunit [112,113].

There is no clear understanding of the mechanisms by which perturbations induced by Omicron mutations alter the conformational landscape of the S protein and how they impact the dynamics of its epitopes and distribution of binding sites including dynamic formation of cryptic binding pockets. In the current study we perform a systematic comparative analysis of the conformational dynamics and allostery in the multiple open and closed functional states of S trimers for BA.1 and BA.2 Omicron variants. By using a combination of coarse-grained and all-atom molecular dynamics (MD) simulations and dynamics network analysis, we identify on large scale the distribution of emerging cryptic binding pockets in the S trimers for BA.1 and BA.2 variants. A network-centric model of screening and ranking the predicted binding pockets from molecular simulations provides an adaptation and extension of the reversed allosteric communication strategy [81–86]. By using this model, we discover commonalties along with significant variant-specific differences in the distribution of cryptic binding sites in the BA.1 and BA.2 trimers, suggesting that small variations could lead to different preferences in allocation of druggable sites. We show that the proposed approach can recover the experimentally known allosteric sites in the NTD, RBD regions as well as targetable S2 regions as highly ranked top binding pockets. The results of this study indicate that while both closed and open S-BA.1 trimers featured as highly ranked NTD supersite region [63], the conformational dynamics of the S-BA.2 trimers could mask the NTD cryptic region and present previously underappreciated cryptic binding pockets in the inter-protomer interface and hinge regions of the S2 subunit. Using this approach, we demonstrate that clusters of allosteric hotspots mediating long-range communications in the S trimers could anchor the druggable cryptic binding pockets, including the experimental allosteric sites. The results of this study are particularly significant for understanding the universe of cryptic bindings sites and variant-specific preferences for druggable pockets. This may have implications for discovery and design of fusion inhibitors and modulators that due to their small size and specific conformation are more likely to bind the cryptic epitopes that are usually inaccessible to conventional antibodies, inducing more potent cross-neutralizing activity.

## 2. Materials and Methods

### 2.1 Structural modeling and refinement

The crystal structures of the S trimers in the open and closed forms for BA.1 and BA.2 variants (Tables 1,2) were obtained from the Protein Data Bank [114]. During structure preparation stage, protein residues in the crystal structures were inspected for missing residues and protons. Hydrogen atoms and missing residues were initially added and assigned according to the WHATIF program web interface [115]. The protonation states for all the titratable residues of the ACE2 and RBD proteins were predicted at pH 7.0 using Propka 3.1 software and web server [116,117]. The missing segments in the studied structures of the SARS-CoV-2 S protein were reconstructed and optimized using template-based loop prediction approach ArchPRED [118].The side chain rotamers were refined and optimized by SCWRL4 tool [119]. The protein structures were then optimized using atomic-level energy minimization with composite physics and knowledge-based force fields implemented in the 3Drefine method [120].

### 2.2 Coarse-Grained Dynamics Simulations

Coarse-grained Brownian dynamics (BD) simulations have been conducted using the ProPHet (Probing Protein Heterogeneity) approach and program [121–124]. BD simulations are based on a high resolution CG protein representation [125] of the SARS-CoV-2 S Omicron trimer structures that can distinguish different residues. In this model, each amino acid is represented by one pseudo-atom at the Cα position, and two pseudo-atoms for large residues. The interactions between the pseudo-atoms are treated according to the standard elastic network model (ENM) in which the pseudo-atoms within the cut-off parameter, *R*_c_ = 9 Å are joined by Gaussian springs with the identical spring constants of *γ* = 0.42 N m^−1^ (0.6 kcal mol^−1^ Å^−2^. The simulations use an implicit solvent representation via the diffusion and random displacement terms and hydrodynamic interactions through the diffusion tensor using the Ermak-McCammon equation of motions and hydrodynamic interactions as described in the original pioneering studies that introduced Brownian dynamics for simulations of proteins [126,127]. The stability of the SARS-CoV-2 S Omicron trimers was monitored in multiple simulations with different time steps and running times. We adopted Δ*t* = 5 fs as a time step for simulations and performed 100 independent BD simulations for each system using 500,000 BD steps at a temperature of 300 K. The CG-BD conformational ensembles were also subjected to all-atom reconstruction using PULCHRA method [128] and CG2AA tool [129] to produce atomistic models of simulation trajectories.

### 2.3 All-Atom Molecular Dynamics Simulations

NAMD 2.13-multicore-CUDA package [130] with CHARMM36 force field [131] was employed to perform 500 ns all-atom MD simulations for representative closed and open S-BA.1 structures (pdb id 7WK2, 7WK3) and closed and open S-BA.2 trimer structures respectively (pdb id 7XIX,7XIW). The structures of the SARS-CoV-2 S-RBD complexes were prepared in Visual Molecular Dynamics (VMD 1.9.3) [1132] by placing them in a TIP3P water box with 20 Å thickness from the protein. Assuming normal charge states of ionizable groups corresponding to pH = 7, sodium (Na+) and chloride (Cl-) counter-ions were added to achieve charge neutrality and a salt concentration of 0.15 M NaCl was maintained. All Na^+^ and Cl^−^ ions were placed at least 8 Å away from any protein atoms and from each other. The long-range non-bonded van der Waals interactions were computed using an atom-based cutoff of 12 Å with the switching function beginning at 10 Å and reaching zero at 14 Å. SHAKE method was used to constrain all bonds associated with hydrogen atoms. Simulations were run using a leap-frog integrator with a 2 fs integration time step. ShakeH algorithm of NAMD was applied for water molecule constraints. The long-range electrostatic interactions were calculated using the particle mesh Ewald method [133] with a cut-off of 1.0 nm and a fourth order (cubic) interpolation. Simulations were performed under NPT ensemble with Langevin thermostat and Nosé-Hoover Langevin piston at 310 K and 1 atm. The damping coefficient (gamma) of the Langevin thermostat was 1/ps. The Langevin piston Nosé-Hoover method in NAMD is a combination of the Nose-Hoover constant pressure method [134] with piston fluctuation control implemented using Langevin dynamics [135,136]. Energy minimization was conducted using the steepest descent method for 100,000 steps. All atoms of the complex were first restrained at their crystal structure positions with a force constant of 10 Kcal mol^−1^ Å^−2^. Equilibration was done in steps by gradually increasing the system temperature in steps of 20K starting from 10K until 310 K and at each step 1ns equilibration was done keeping a restraint of 10 Kcal mol-1 Å-2 on the protein C_α_ atoms. After the restraints on the protein atoms were removed, the system was equilibrated for an additional 10 ns. An NPT production simulation was run on equilibrated structures for 500 ns keeping the temperature at 310 K and constant pressure (1 atm).

### 2.4 Network Analysis of Conformational Ensembles

A graph-based representation of protein structures [137,138] is used to represent residues as network nodes and the inter-residue edges to describe non-covalent residue interactions. The weights of the network edges in the residue interaction networks are determined by dynamic residue cross-correlations obtained from MD simulations [139] and coevolutionary couplings between residues measured by the mutual information scores [140]. Residue Interaction Network Generator (RING) program [141] was employed for generation of the initial residue interaction networks. The edge lengths in the network are then adjusted using the generalized correlation coefficients associated with the dynamic correlation and mutual information shared by each pair of residues. Network edges are weighted for residue pairs within at least one independent simulation. Network analysis was performed using the python package NetworkX [142]. The short path betweenness of residue *i* is defined to be the sum of the fraction of shortest paths between all pairs of residues that pass through residue *i*:

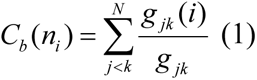

Where *g _jk_* denotes the number of shortest geodesics paths connecting *j* and *k,* and *g _jk_* (*i*) is the number of shortest paths between residues *j* and *k* passing through the node *n_i_.* Residues with high occurrence in the shortest paths connecting all residue pairs have a higher betweenness values. For each node *n,* the betweenness value is normalized by the number of node pairs excluding *n* given as (*N* −1)(*N* - 2) / 2, where *N* is the total number of nodes in the connected component that node *n* belongs to.

The Girvan-Newman algorithm [143] is used to identify local communities. An improvement of Girvan-Newman method was implemented where all highest betweenness edges are removed at each step of the protocol. The algorithmic details of this modified scheme were presented in our recent study [144]. A community-based model of allosteric communications is based on the notion that groups of residues that form local interacting communities are expected to be highly correlated and can switch their conformational states cooperatively. In this model, long-range allosteric communications can be transmitted not between individual residue nodes but rather through a hierarchical chain of local communities connected via inter-modular bridges on the dynamic interaction networks. Using the local community decomposition in the protein structures, we identify a hierarchy of clusters consisting of dynamically and coevolutionary coupled nodes (residues at the hierarchical level 1 and meta-nodes formed by small clusters at the next hierarchical level 2). To determine inter-modular bridges between local communities, we employed the inter-community centrality metric [145]. This parameter uses community detection as input :

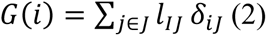

where the sum is over communities *J* (different from the community of node *i*, denoted as *I*), δ_*iJ*_ is equal 1 if there is a link between node *i* and community *J* and 0 otherwise. *l*_*iJ*_ corresponds to the distance between community *I* and community *J*. It is measured by the inverse of the number of links between them. The community centrality is determined according to the following expression :

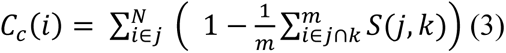

where N is a number of communities to which node i belongs, and S the Jaccard similarity coefficient between community j and k, calculated for the number of shared nodes between each community pair. The sum is averaged over the m communities that are paired with community j to which a given node i may simultaneously belong. All parameters were computed using the python package NetworkX [142].

### 2.5 Machine Learning-Based Discovery of Cryptic Pockets and Network-Based Ranking of Allosteric Pocket Propensities and Allosteric Binding Sites

We used two different complementary approaches for enumerating the putative binding pockets in the conformational ensembles of the S-BA1 and S-BA.2 structures. A template-free P2Rank approach is among the most efficient and fast available ML tools for prediction of ligand binding sites that combines sequence and structural data to rank potential binding sites based on their likelihood of binding a specific ligand [103,104]. P2Rank uses support vector machine (SVM), random forests (RF), and artificial neural networks (ANNs) to learn the ligandability of a local chemical environment that is centered on points placed on the protein’s solvent-accessible surface [103,104]. P2Rank v2.4 with default parameters was deployed to identify pockets across all of the representative states from our simulations. By combining eXtreme gradient boosting (XGBoost) and graph convolutional neural networks (GCNNs) a robust approach for allosteric site identification and Prediction of Allosteric Sites Server (PASSer) was developed [106–108]. We also employed the PASSer Learning to Rank (LTR) model that is capable of ranking pockets in order of their likelihood to be allosteric sites. In this approach, FPocket was applied on each protein to detect protein pockets [146]. For each detected pocket, 19 physical and chemical features are calculated, ranging from pocket volume, solvent accessible surface area to hydrophobicity. The LTR model uses LightGBM [147] one of the two popular implementations of Gradient-boosted decision trees (GBDT) method and XGBoost [148]. Using P2Rank [103,104] and PASSer LTR [107] approaches, we identified binding pockets in the conformational ensembles and computed P2Rank-predicted residue pocket probability. In the next step, we reweighted the pocket propensities scores and re-ranked the predicted binding pockets, using network-based hierarchical community centralities determined for each protein residue. We then characterize the difference in each residue’s maximum ligand-binding pocket probability in the ensemble and network-weighted pocket propensities. The reported top binding pockets for each protein structure correspond to network-ranked consensus P2Rank/LTR predicted sites. These binding pockets are formed by residues with high network centrality scores that mediate allosteric interactions in the protein structures. We refer to these cryptic binding pockets as potential allosteric binding sites.

## 3. Results

### 3.1. Conformational Landscapes of Multiple Conformational States of the SARS-CoV-2 S BA.1 and BA.2 Trimers

We combined multiple coarse-grained simulations followed by atomistic reconstruction of the trajectories to perform a comparative analysis of the conformational landscapes for S Omicron BA.1 (Figure 1) and BA.2 trimers (Figure 2). A total of 12 Omicron mutations (G339D, S373P, S375F, K417N, N440K, S477N, T478K, E484A, Q493R, Q498R, N501Y, and Y505H) are shared among the BA.1 and BA.2 variants. In the RBD, BA.1 contains unique mutations S371L, G446S, and G496S while BA.2 carries S371F, T376A, D405N, and R408S mutations (Table 1). The BA.1 and BA.2 S trimers are different in three additional residues, T547K in the S1 region, D856K and L981F in S2 region that are found in the BA.1 but not in the BA.2 variant (Table 1). In the conformational dynamics analysis, we simulated all available cryo-EM structures of the unbound S-BA.1 and S-BA.2 trimers in the 3 RBD-down closed form, various 1RBD-up open conformations and also 2RBD-up state. Using coarse-grained Brownian dynamics (CG-BD) within the ProPHet (Probing Protein Heterogeneity) approach [121–124], we performed 100 independent simulations for each structure (Table 2) to produce sufficient sampling. These simulations were also complemented by atomistic MD simulations performed for representative closed and open S-BA.1 (pdb id 7WK2, 7WK3) and closed and open S-BA.2 trimer structures (pdb id 7XIX,7XIW) (Table 3). Multiple structures of the S Omicron trimers used in CG-BD and MD simulations (Tables 2,3) allowed for a comparative analysis of protein dynamics in the distinct S Omicron states.

**Figure 1.**
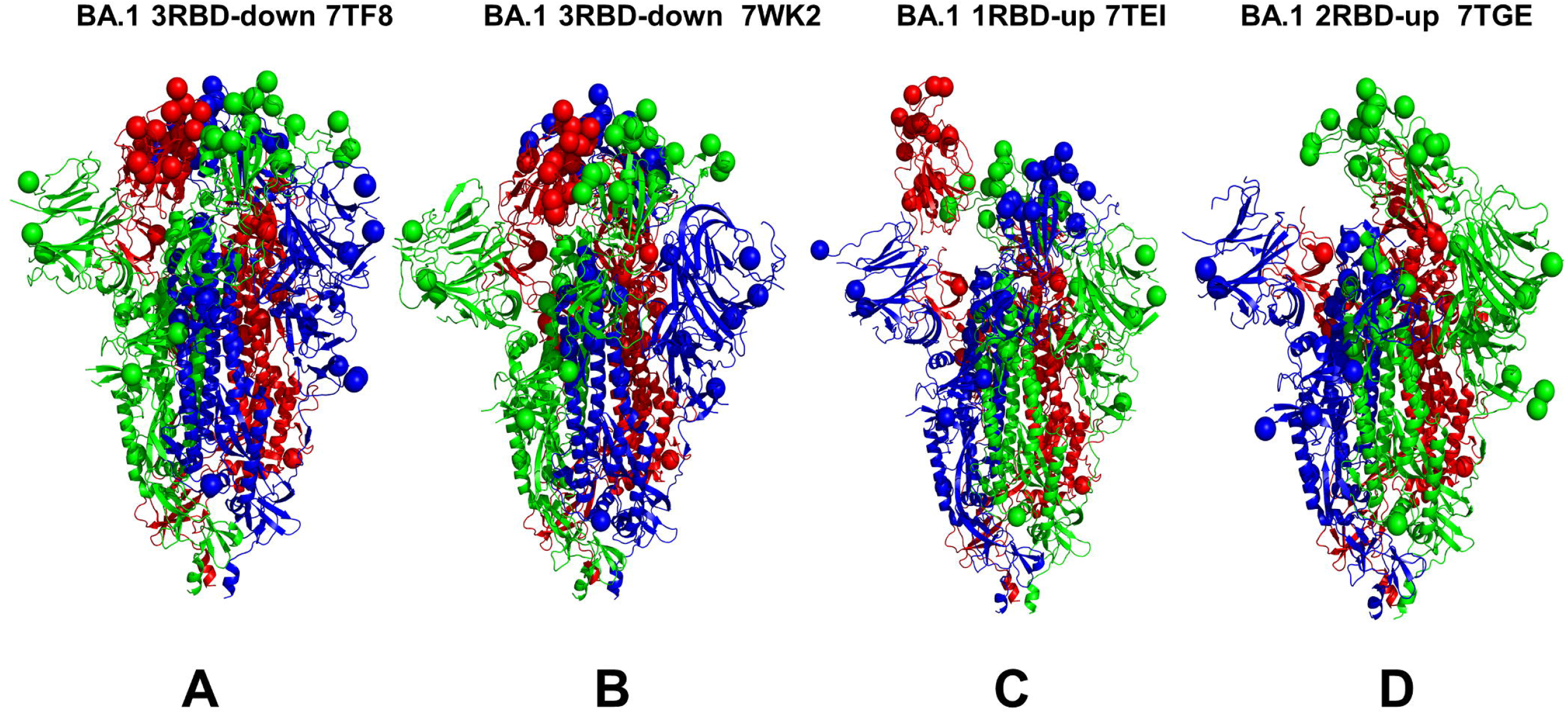
Structural organization and mapping of the Omicron BA.1 mutations in the SARS-CoV-2 S closed Omicron BA.1 trimers. A general overview of the SARS-CoV-2 S Omicron BA.1 trimer in the 3RBD-down closed form, pdb id 7TF8 (A), BA.1 closed trimer, pdb id 7WK2 (B), BA.1 1RBD-up open trimer, pdb id 7TEI (C) and BA.1 2RBD-up open trimer, pdb id 7TGE (D). Protomer A is shown in green ribbons, protomer B is in red ribbons, and protomer C is in blue ribbons. The positions of the Omicron mutations are shown for each protomer in spheres colored according to the respective protomer.

**Figure 2.**
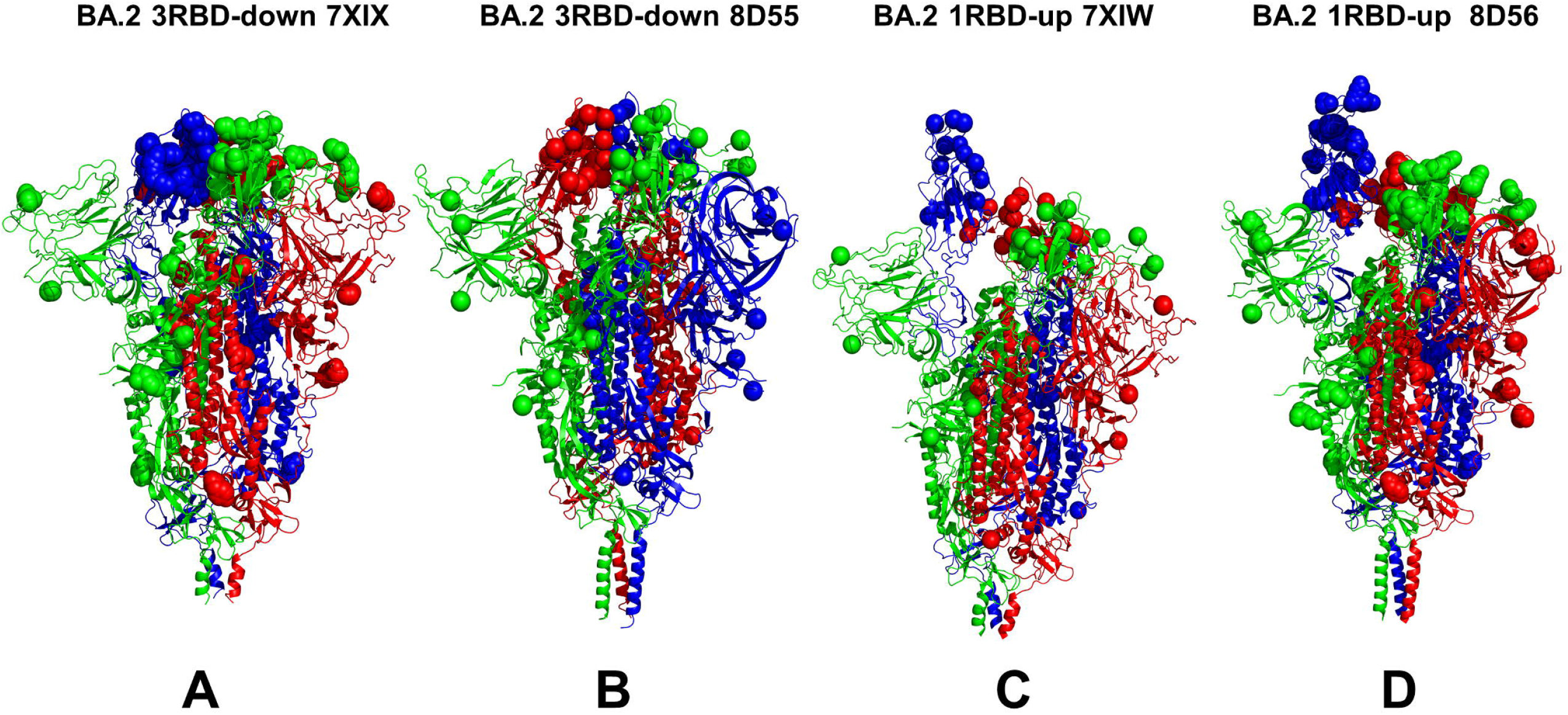
Structural organization and mapping of the Omicron BA.2 mutations in the SARS-CoV-2 S closed Omicron BA.2 trimers. A general overview of the SARS-CoV-2 S Omicron BA.2 trimer in the 3RBD-down closed form, pdb id 7XIX (A), BA.2 closed trimer, pdb id 8D55 (B), BA.2 1RBD-up open trimer, pdb id 7XIW (C) and BA.2 1RBD-up open trimer, pdb id 8D56 (D). Protomer A is shown in green ribbons, protomer B is in red ribbons, and protomer C is in blue ribbons. The positions of the Omicron BA.2 mutations The Omicron BA.2 mutational sites are shown for each protomer in spheres colored according to the respective protomer.

**Table 1.** Mutational landscape of the Omicron BA.1 and BA.2 mutations.

| Omicron Variant | Mutational landscape |
| --- | --- |
| BA.1 | A67, T95I, G339D, S371L, S373P, S375F, K417N, N440K, G446S, S477N, T478K, E484A, Q493R, G496S, Q498R, N501Y, Y505H, T547K, D614G, H655Y, N679K, P681H, N764K, D796Y, N856K, Q954H, N969K, L981F |
| BA.2 | T19I, G142D, V213G, G339D, S371F, S373P, S375F, T376A, D405N, R408S, K417N, N440K, S477N, T478K, E484A, Q493R, Q498R, N501Y, Y505H, D614G, H655Y, N679K, P681H, N764K, D796Y, Q954H, N969K |

**Table 2.** Structures of SARS-CoV-2 S Omicron protein structures examined in this study.

|  | **PDB**<br>**code** | **Description** | **RBDs position** |
| --- | --- | --- | --- |
| BA.1 | 7TF8 | SARS-CoV- 2 S Omicron Trimer, closed | 3 RBD-down |
| BA.1 | 7WK2 | SARS-CoV- 2 S Omicron Trimer, closed | 3RBD-down |
| BA.1 | 7TNW | SARS-CoV- 2 S Omicron Trimer, closed | 3 RBD- down |
| BA.1 | 7TL1 | SARS-CoV- 2 S Omicron Trimer, closed | 3 RBD-down |
| BA.1 | 7TL9 | SARS-CoV- 2 S Omicron Trimer, open | 1RBD-up |
| BA.1 | 7TEI | SARS-CoV- 2 S Omicron Trimer, open | 1 RBD-up |
| BA.1 | 7WK3 | SARS-CoV- 2 S Omicron Trimer, open | 1RBD-up |
| BA.1 | 7TO4 | SARS-CoV- 2 S Omicron Trimer, open | 1RBD-up |
| BA.1 | 7WVN | SARS-CoV- 2 S Omicron Trimer, open | 1RBD-up |
| BA.1 | 7WVO | SARS-CoV- 2 S Omicron Trimer, open | 1RBD-up |
| BA.1 | 7TGE | SARS-CoV- 2 S Omicron Trimer, open | 2 RBD-up |
| BA.2 | 7XIX | SARS-CoV- 2 S Omicron Trimer, closed | 3 RBD-down |
| BA.2 | 7UB0 | SARS-CoV- 2 S Omicron Trimer, closed | 3 RBD-down |
| BA.2 | 7UB5 | SARS-CoV- 2 S Omicron Trimer, closed | 3 RBD-down |
| BA.2 | 7UB6 | SARS-CoV- 2 S Omicron Trimer, closed | 3 RBD-down |
| BA.2 | 8D55 | SARS-CoV- 2 S Omicron Trimer, closed | 3 RBD-down |
| BA.2 | 7XIW | SARS-CoV- 2 S Omicron Trimer, open | 1RBD-up |
| BA.2 | 8D56 | SARS-CoV- 2 S Omicron Trimer, open | 1RBD-up |

**Table 3.** Molecular simulations of the RBD-ACE2 complexes.

| **PDB** | **System** | **CG-BD** | **#simulations** | **All-atom MD** |
| --- | --- | --- | --- | --- |
| 7WK2 | S-BA.1 3-RBD down | 500,000<br>steps | 100 | 500 ns |
| 7WK3 | S-BA.1 1-RBD up | 500,000<br>steps | 100 | 500 ns |
| 7XIX | S-BA.2 3-RBD down | 500,000<br>steps | 100 | 500 ns |
| 7XIW | S-BA.2 1-RBD up | 500,000<br>steps | 100 | 500 ns |

Conformational dynamics of the S-BA.1 and S-BA.2 closed conformations showed generally similar root-mean-square-fluctuation (RMSF) profiles reflecting stability of the 3-RBD structure for both variants (Figure 3A,B). The profiles showed similar thermal fluctuations in the RBD-closed S1 subunit (residues 14-530) and a more rigid S2 subunit (Figure 3A,B). Greater deviations were detected in the NTD (residues 14-306) but the pattern of thermal fluctuations in this region differs between BA.1 and BA.2 trimers. In the S-BA.1 trimer we observed larger RMSF displacements in the NTD residues 75-250 (Figure 2A), while for the S-BA.2 variant we observed several NTD peaks corresponding to residues 60-80, 140-180 and 240-260 (Figure 3B). In the Omicron BA.1 and BA.2 closed trimers, the upstream helix (UH) (residues 736-781) and central helix (CH) (residues 986-1035) are particularly rigid, while C-terminal domain 1, CTD1 (residues 528-591) and C-terminal domain 2, CTD2 (residues 592-686) undergo moderate fluctuations (Figure 3A,B). For both variants, an appreciable degree of flexibility was also seen in the RBD residues 466-498 and residues 615-640 of the CTD2 as well as in the furin cleavage region (residues 670-690) (Figure 3). Consistent with strong sequence conservation, the CH region is exceedingly rigid in the S-BA.1 and S-BA.2 closed trimers. Interestingly, the conformational dynamics showed signs of greater rigidity in the S2 subunit, but more flexibility in the S1 subunit (particularly in the NTD) for the S-BA.1 closed trimer (Figure 3A). At the same time, the RMSF profiles indicated a moderately greater plasticity of both S1 and S2 subunits for the BA.2 variant (Figure 3B). The inter-protomer salt bridges K856-D571 in the BA.1 timers and absence of these stabilizing interactions in the BA.2 trimer may contribute to some plasticity in the S1-S2 and S2 regions for the S-BA.2 trimer.

**Figure 3.**
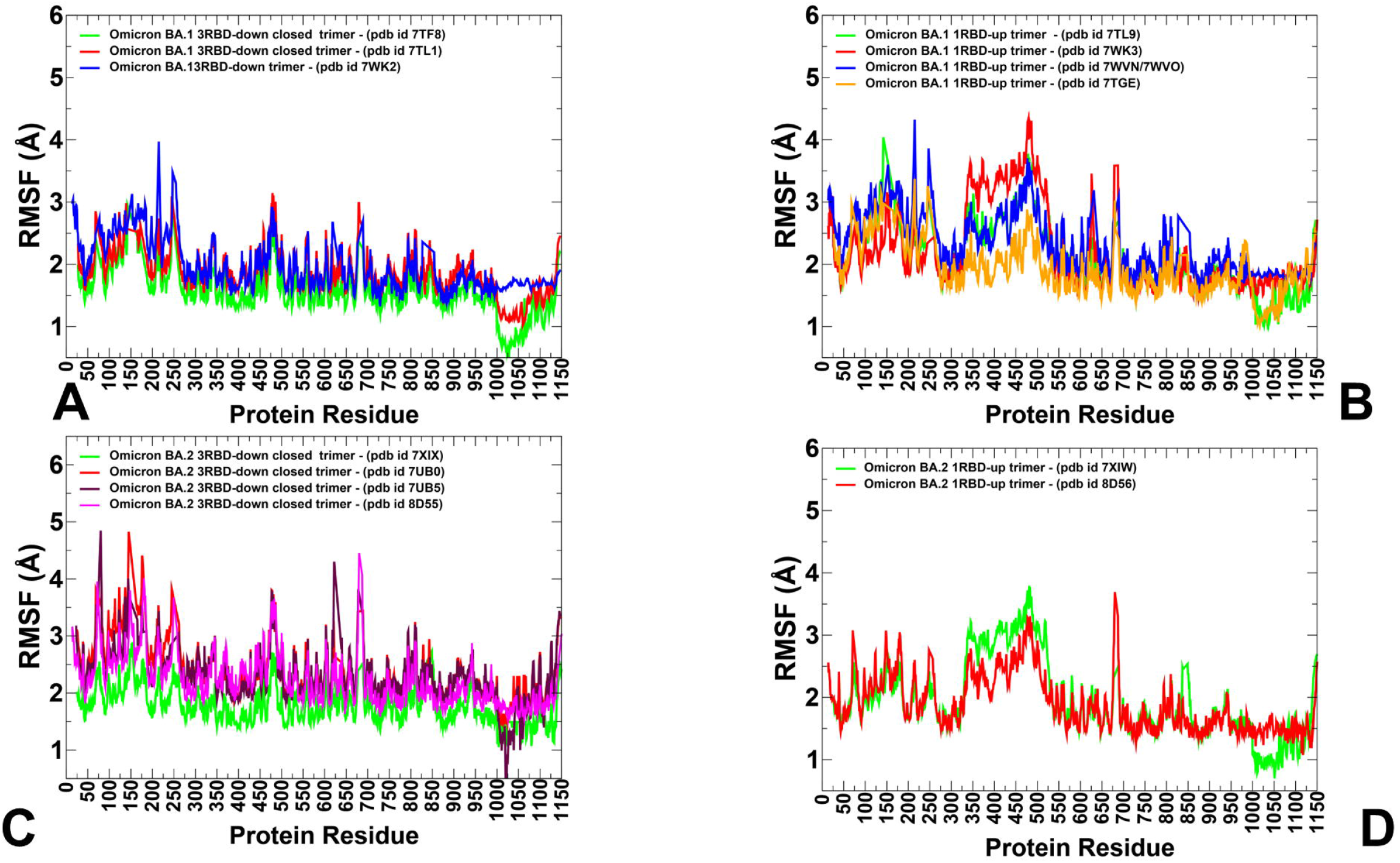
Conformational dynamics profiles obtained from CG-BD and MD simulations of the Omicron BA.1 and BA.2 trimers. (A) The RMSF profiles for the S protein residues obtained from simulations of the S-BA.1 closed trimers (pdb id 7TF8 in green lines, pdb id 7TL1 in red lines, and pdb id 7WK2 in blue lines). (B) The RMSF profiles for the S protein residues obtained from simulations of the S-BA.1 1 RBD-up open trimers (pdb id 7TL9 in green lines, pdb id 7WK3 in red lines, pdb id 7WVN/7WVO in blue lines), and S-BA.1 2RBD-up open trimer (pdb id 7TGE in orange lines). (C) The RMSF profiles for the S protein residues obtained from simulations of the S-BA.2 closed trimers (pdb id 7XIX in green lines, pdb id 7UB0 in red lines, pdb id 7UB5 in maroon lines, and pdb id 8D55 in magenta lines). (D) The RMSF profiles for the S protein residues obtained from simulations of the S-BA.2 1 RBD-up open trimers (pdb id 7XIW in green lines and pdb id 8D56 in red lines).

However, the differences in the RMSF profiles of the closed trimer for the BA.1 and BA.2 variants remained moderate, suggesting thermal stability of the closed states and only marginally increased mobility of the S-BA.2 trimers. A comparison of conformational dynamics profiles obtained from simulations of RBD-up open states for S-BA.1 and S-BA2 trimers revealed several interesting patterns (Figure 3C,D). It was observed that the RMSF distributions for the open S-BA.1 structures showed a larger degree of heterogeneity and greater fluctuations of the NTD and exposed RBD-up regions (Figure 3C). In some contrast, the conformational flexibility of the NTD and RBD in the open BA.2 trimers is moderately reduced, displaying smaller displacements within RMSF < 3 Å (Figure 3D). Based on these fundings, we argue that the RBD-up conformations for the S-BA.1 trimer may experience greater variability in the S1 subunit, while the open states for the S-BA.2 trimer become more stable, thus suggesting that the open S form could become more dominant in the BA.2 variant. These findings are consistent with the experiments showing the increased RBD stability and enhanced propensity to populate open trimer states for the BA.2 variant [46].

In addition to variability of the RBD-up regions, we observed differences in the NTD mobility. In this context, it is worth noting that NTD region (residues 14–20, NTD N-terminus), N3 (residues 141–156), and N5 (residues 246–260) collectively form an antigenic supersite on the NTD. These regions showed the increased RMSF values for both variants, but the degree of conformational heterogeneity appeared to be greater in the S-BA.2 conformations (Figure 3). This suggests that conformationally flexible NTD regions in the S-BA.1 trimers may preserve the topology of the NTD supersite, while specific NTD mutations and the increased heterogeneity of the NTD regions in the BA.2 variant could promote reconstruction of the antibody NTD site and lead to formation of dynamic cryptic pockets on the NTD surfaces. These dynamic characteristics gleaned from RMSF analysis are consistent with thermal stability assays that verified that S BA.2 trimers are the least stable among BA.1, BA.2, BA.3 and BA.4 variants [48].

### 3.2 Functional Dynamics of the BA.1 and BA.2 Trimers Highlights Complementary Roles of the Omicron Mutational Sites as Hinges and Transmitters of Collective Motions

To characterize collective motions in the SARS-CoV-2 S Omicron structures, we performed principal component analysis (PCA) of trajectories. The low-frequency modes can be exploited by mutations to modulate protein movements that are important for allosteric transformations between the open and closed states. The local minima along the slow mode profiles are typically aligned with the hinge centers, while the maxima could point to moving regions undergoing concerted movements. Overall, the key functional signature of collective dynamics in these states are the large displacements of the NTD and RBD regions that enable S protein to assume the receptor-accessible conformation (Figure 4). The important finding of this analysis is the correspondence of the major hinge positions with the Omicron sites in the S2 subunit (Figure 3). The slow mode profiles in the closed S-BA.1 states showed that residues N764K, N856K, Q954H, N969K, and L981F corresponded to the local minima of the distribution (Figure 4A-C). Interestingly, the key immobilized hinge positions are conserved and shared by both closed and open forms of the S-BA.1 and S-BA.2 trimers. A comparison of slow modes for the closed forms of the S-BA.1 (Figure 4A) and S-BA2 trimers (Figure 4D) indicated interesting differences. In the S-BA.1 closed trimer, not only the closed-down RBDs are mostly immobilized in collective motions but also the NTD regions may experience only moderate functional displacements (Figure 4A). On the other hand, functional movements of the NTD regions are more pronounced and diverse among the three protomers in the S-BA.2 closed trimer (Figure 4D). These observations suggested a greater heterogeneity of functional NTD movements in the BA.2 closed state. The results are consistent with the conformational dynamics profiles showing the increased plasticity of the NTD regions in the otherwise stable S-BA.2 closed trimers. As may be expected, in the open BA.1 conformations we observed significant functional displacements of the RBD region, indicating that Omicron mutations target sites involved in collective movements (Figure 4B,C). Consistent with our previous studies of the S trimer protein, the functional dynamics profiles also revealed that F318, A570, I572 and F592 residues are conserved hinge sites that are situated near the inter-domain CTD1-S2 interfaces and could function as regulatory centers governing functional transitions (Figure 4B,C). There are differences between the slow mode profiles for the open trimer states of the BA.1 and BA.2 variants. For the S-BA.1 open trimer the profiles showed significant displacements of both NTD and RBD regions (Figure 4B,C). Moreover, the functional NTD motions can occur in both closed and open protomers of the 1RBD-up BA.1 trimer, leading to significant variations of these regions, while the S2 subunit is immobilized. As a result, the S-BA.1 trimers may be prone to conformational transformations between the open and closed forms in the absence of the host receptor, where ACE2 binding promotes shift to the open form.

**Figure 4.**
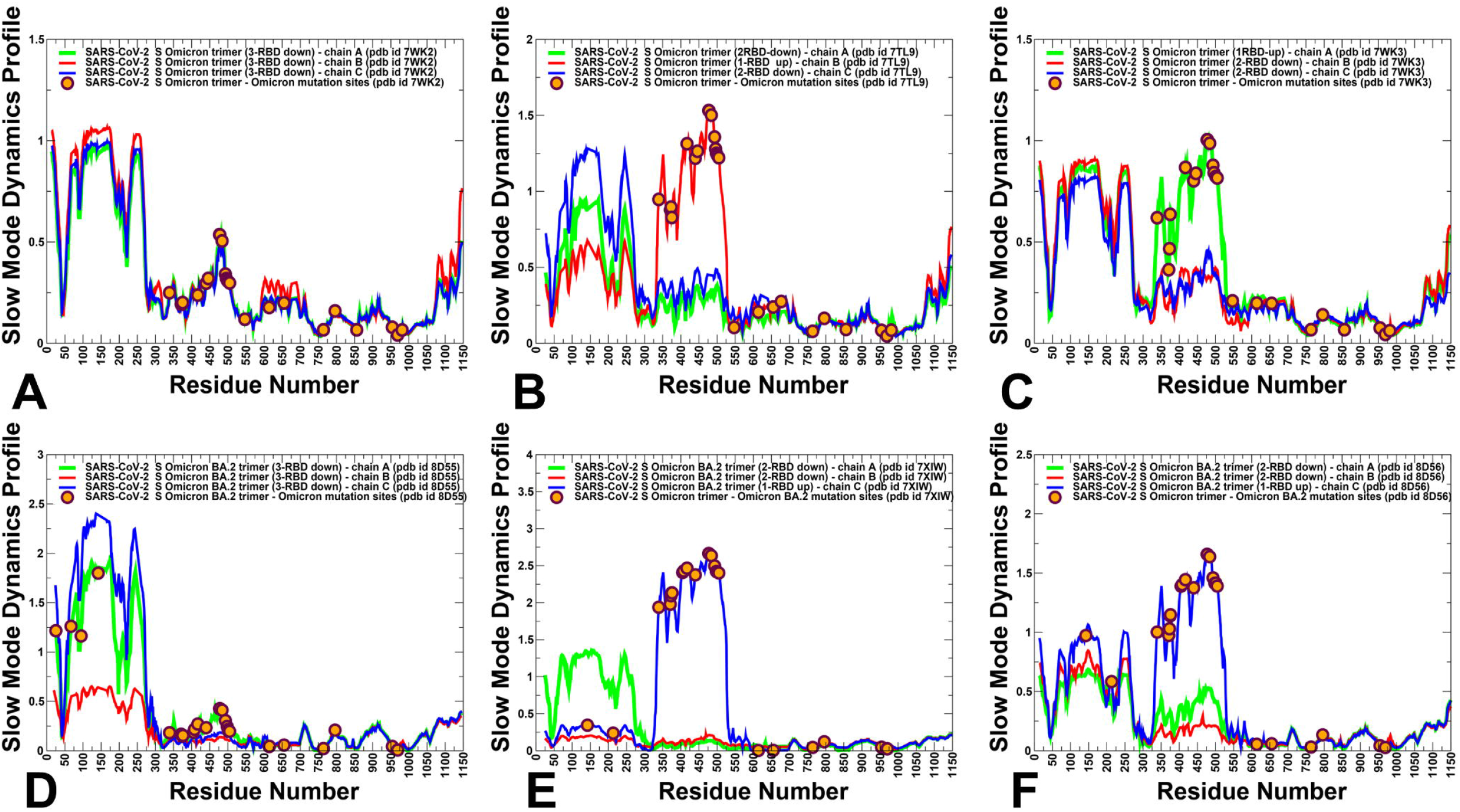
The slow mode mobility profiles of the SARS-CoV-2 S Omicron trimer structures. The slow mode dynamics profiles represent the displacements along slow mode eigenvectors and correspond to the cumulative contribution of the slowest five modes. The slow mode mobility profiles for the S-BA.1 trimer states : 3RBD-down closed BA.1 trimer, pdb id 7WK2 (A), 1RBD-up open BA.1 trimer, pdb id 7TL9 (B), and 1RBD-up open BA.1 trimer, pdb id 7WK3 (C). The slow mode mobility profiles for the S-BA.2 trimer states: 3RBD-down closed BA.2 trimer, pdb id 8D55 (D), 1RBD-up open BA.2 trimer, pdb id 7XIW (E), and 1RBD-up open BA.2 trimer, pdb id 8D56 (F). The slow mode profiles for protomer chains A, B and C are shown in green, red and blue lines respectively. The positions of the Omicron BA.1 and BA.2 mutational sites are shown in orange-colored filled circles.

In contrast, for the S-BA.2 open states, the movements of the S1 subunit become mainly associated with the RBD movements while NTD displacements are less significant (Figure 4E,F). Accordingly, while the intrinsic heterogeneity of the NTD regions is manifested in significant collective movements of the closed S-BA.2 conformations, functional displacements of the NTD regions become reduced in the open S-BA.2 form signaling stabilization of the open state. This indicates that the 1RBD-up open trimer conformation may be a preferable state for the S-BA.2 variant, leading to a greater propensity of the BA.2 variant for binding with the host receptor. This analysis supports a mechanism in which dynamic preferences for the S-BA.2 open forms and more RBD-up conformations may cause higher transmission and infection rates of the BA.2 Omicron sublineage compared to BA.1.

The observations from the functional dynamics analysis are supported by the findings from thermal shift assays showing the observed lower temperature for dissociation of the S-BA.2 trimer [46]. Moreover, this experimental study showed the dissociation constant of the BA.2 S protein with ACE2 is fivefold and twofold higher than that of the wild type S protein and S-BA.1 protein with ACE2, respectively [46]. Our analysis of collective movements suggests that evolution of the Omicron variants may be accompanied by subtle changes in the intrinsic dynamic patterns leading to progressive shift towards more stable open form for the BA.2 variant. The observed patterns of collective motions in the S-BA.1 and S-BA.2 states are also consistent with the recent experiments showing the increasing flexibility of the NTD regions in the Omicron trimers which may enable escape of NTD-targeting antibodies thereby providing an evolutionary advantage. The observed in our dynamics analysis differences in modulation of the NTD mobility between BA.1 and BA.2 conformations supports the proposed mechanism in which NTD provides an adaptable antigenic surface, allowing for enhanced immune escape potential [50].

Interestingly, several Omicron sites belong to local hinge regions (N764K, D796Y, N856K, Q954H, N969K, and L981F), while RBD mutational sites are located in regions prone to functional movements. The hinge sites F318, A570, T572, F592, D614G, N764K, N856K, Q954H, and N969K. participate in direct interaction contacts that can strengthen stability of the hinge clusters in the S-BA.1 trimers. In particular, N856K interacts with T572, while N764K makes favorable contacts T315, N317 and Q314 positions of the adjacent protomer. N969K is hydrogen-bonded with Y756 and Q755 sites, while T547K is in direct contacts with S982 and L981F positions. The Omicron mutations in these regions introduce new stabilizing contacts that promote structural rigidity and expansion of the hinge clusters. We also noticed that F318, S383, A570, I572 and F592, D985 residues are conserved hinge sites that are situated near the inter-domain SD1-S2 interfaces and could function as regulatory centers governing the inter-protomer and inter-domain transitions (Figure 4). In both the closed and open forms of the S Omicron variants the NTD Omicron mutational sites experience large displacements and allow for remodeling of the NTD binding epitope thus reducing the effect of antibody-mediated protection. At the same time, the RBD Omicron sites in the open state belong to highly mobile in collective dynamics regions (Figure 4). Given that Omicron mutational positions in the S2 subunit tend to occupy immobilized hinge sites, this suggests long-range communications and allosteric cross-talk between S2 and S1 subunits may be orchestrated and modulated by the Omicron mutations. In this context, it may be also argued that the increased dynamic preferences of BA.2 for the open states with flexible NTD and RBD regions may promote binding with ACE2 while leveraging the adaptability of the NTD binding surfaces to escape immunity.

### 3.3 Dynamic Network Analysis: Variant-Induced Modulation of Allosteric Mediating Centers

We used hierarchical network modeling to characterize the organization of residue interaction networks in the S Omicron functional states. We constructed and analyzed the dynamic residue interaction networks using an approach in which the dynamic fluctuations obtained from simulations are mapped onto a graph with nodes representing residues and edges representing weights of the measured dynamic properties. The hierarchical community centrality metric is employed in this network model to identify S protein positions that mediate allosteric communications between local communities (Figure 5). As a result, residues featuring high community centrality values may function as inter-community connectors. For clarity of presentation and consistency, we report the distributions of community centrality where the inter-community bridges are represented by at the single residue level. We found that in the BA.1 closed trimers, the distinct maxima of the network distribution correspond to the clusters of residues in the NTD (residues 50-60, 100-120, 200-207) (Figure 5A). Strikingly, some of these NTD residues contribute to the dynamically evolving binding pocket formed by residues I101, W104, I119, V126, M177, F192, F194, I203, and L226 which form the interactions with the ligand in the cryo-EM structure [61,62]. These NTD peaks are mostly preserved in the open S-BA.1 trimers (Figure 5B) but the contributions of these regions to the network distribution is modestly reduced. Importantly, we found that for all protomers in the closed state, the dominant centrality peak corresponds to the N2R linker region (residues 300-336) that connects the NTD and RBD within a single protomer unit. CTD1 region and strong centrality density in the NTD regions positions (Figure 4A,B). In the S-BA.1 trimers, we also noticed a significant peak associated with the RBD-CTD1 interfaces that harbor stable local communities of residues (C-535 C-554 C-583; A-544 A-564 A-579; B-330 B-544 B-579; C-544 C-564 C-579). The important mediating role of the inter-protomer interfaces is exemplified by the peaks associated with the bridging communities linking neighboring protomers (B-855 C-589 C-592; A-568 A-574 C-854; A-740 A-857 B-592; B-775 B-864 C-665; A-592 C-737 C-855). A broad region of high centrality regions in the closed S-BA.1 trimer corresponds to CTD1 (residues 528-591) and CTD2 (residues 592-686) that links RBD to the S2 regions (Figure 5A). These peaks are weakened in the open BA.1 trimer, thus highlighting changes in allosteric communications between the closed and open forms (Figure 5B). The maxima of the distribution profile are associated with the residues involved in the inter-protomer bridges N764K-Q314, S982-T547K, N856K-D568, N856K-T572, N969K-Q755, N969K-Y756, S383-D985, F855-G614, V963-A570, N317-D737, R319-D737, R319-T739, R319-D745, and K386-F981. The dense cluster peaks of the network profile in the S2 regions correspond to stable UH region (residues 736-781) and CH (residues 986-1035) in which the network centrality peaks often align with the hinge regions (residues 569-572, 756-760). A very pronounced centrality peak is aligned with the HR1/HR2 linker in the S2 subunit (residues 990-1050) that connects HR1 (residues 910-985) and HR2 (residues 1163–1206) (Figure 5,B). The local communities in the HR1/HR2 region (C-1083 C-1088 C-1137; B-1032 B-1043 B-1048; B-1032 B-1048 B-1051; A-1028 A-1043 A-1064; A-1005 B-1005 C-1005) enable stabilization of the S2 regions.

**Figure 5.**
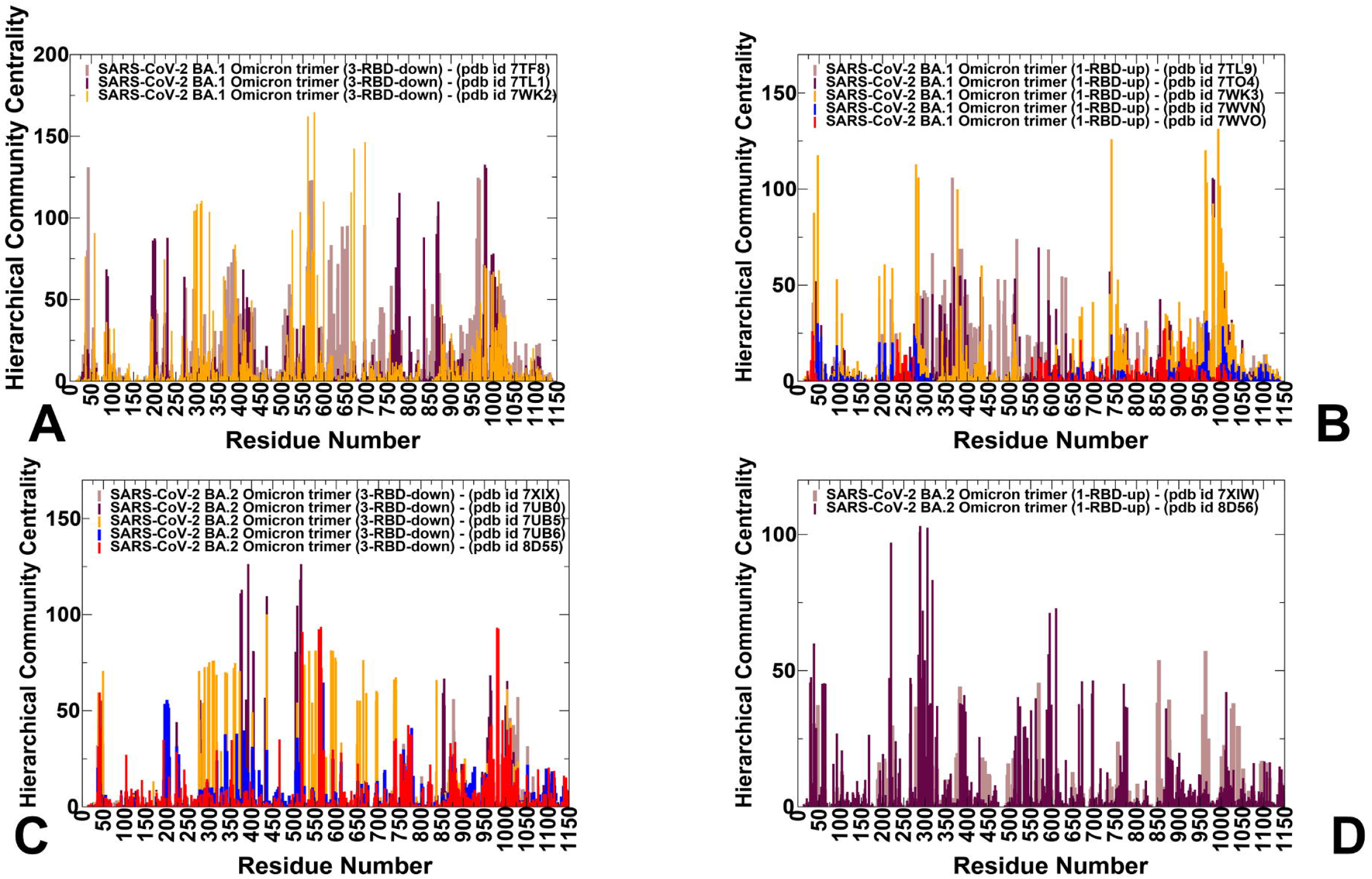
The hierarchical community centrality distributions for the SARS-CoV-2 S Omicron BA.1 and BA.2 structures. (A) The centrality profiles for the for the closed S-BA.1 trimers (pdb id 7TF8 in brown, pdb id 7TL1 in maroon and pdb id 7WK2 in orange-colored bars). (B) The centrality profiles for the for the open S-BA.1 trimers (pdb id 7TL9 in brown, pdb id 7TO4 in maroon, pdb id 7WK3 in orange, pdb id 7WVN in magenta and pdb id 7WVO in red bars). (C) The centrality profiles for the for the closed S-BA.2 trimers (pdb id 7XIX in brown, pdb id 7UB0 in maroon, pdb id 7UB5 in orange, pdb id 7UB6 in magenta, and pdb id 8D55 in red bars). (D) The centrality profiles for the for the open S-BA.2 trimers (pdb id 7XIW in brown and pdb id 78D56 in maroon bars).

We observed some noticeable weakening of the network centrality profiles in the S-BA.2 trimers (Figure 5C,D) which signals a more dynamic and diffuse nature of the interactions for BA.2. The contribution of the NTD regions is reduced as compared to the other profile peaks. At the same time, in the closed S-BA.2 trimer, the largest peak is shifted towards the NTD-RBD and RBD regions as evidenced by high centrality values for residues 350-450 and 500-530 of the RBD (Figure 5C). The RBD-CTD1 interface resides and HR1-HR2 regions also contributed significantly to the distribution. The profile of the open S-BA.2 trimer is less dense suggesting the increased heterogeneity of the long-range communications, while preserving topologically important mediating clusters in the NTD, RBD and CTD1 regions (Figure 5D). Overall, the analysis confirmed the hypothesis that S-BA.2 trimers may exhibit greater conformational plasticity and dynamic changes, which may suggest more room for opening and closing cryptic binding pockets in the S1 and S2 subunits. We also propose that the distribution of the potential allosteric pockets in the S-BA.2 trimers may be more sensitive to the specific structure and display more heterogeneous pocket landscape, while greater structural stability of the S-BA.1 trimers may lead to limited repertoire of dynamically adaptable allosteric sites.

### 3.4 Allostery-Guided Network Screening of Cryptic Binding Pockets in the S-BA.1 and S-BA.2 Conformational Ensembles : Variant-Specific Modulation of the NTD Binding Sites

We employed the conformational ensembles of the S-BA.1 and S-BA.2 trimers in combination with a template-free and highly efficient P2Rank and PASSerRank approaches to describe the available spectrum of potential cryptic sites and compute the residue-based pocket propensities in the ensembles of the S proteins. We combined these robust tools for enumeration of potential pockets with a network-based weighing of pocket probabilities to enable pocket ranking based on allosteric function which provides an adaptation and extension of the reversed allosteric communication strategy [81–86]. The central result of this analysis is discovery of variant-specific differences in the distribution of cryptic binding sites in the BA.1 and BA.2 trimers, suggesting that small variations could lead to different preferences in allocation of druggable sites (Figures 6,7). First, we determined potential binding pockets in the conformational ensembles and evaluated the ligand-binding pocket propensities for each protein residue. We propose that by using network-based hierarchical community centrality parameter as an allostery-based weighting factor, we can adjust the binding pocket residue propensities and provide ranking of the cryptic binding pockets emerging in the conformational ensembles of the S Omicron trimers. By identifying local residue clusters that correspond to the major peaks of the network centrality distributions, we infer the positions of regions that could overlap with potential cryptic binding pockets involved in mediating long-range interactions and communications (Figures 6,7). In our model, we assumed that the predicted binding pockets that overlap and harbor high network centrality sites may correspond to functionally relevant allosteric sites and allow for effective screening/ranking of the identified cryptic pockets. Accordingly, the residue-based binding pocket probability in the ensemble obtained from P2Rank and PASSerRank calculations are weighted based on the residue network centrality values. The reported pocket propensity scores represent ensemble-averaged values, and pockets that are persistent in the equilibrium trajectories are reported (Figures 6,7). We first analyzed the binding pocket propensity scores for the closed and open S-BA.1 trimer conformations (Figure 6). The central finding of this analysis is a surprising emergence of the highest distribution peaks aligned with the NTD regions in the closed S-BA.1 trimers (Figure 6A-C). Consistently, the NTD regions (including residues I101, R102, W104, I119, N121, V126, I128, S172, R190, F192, F194, I203, H207, L226, D227, S94, N99) corresponded to the highest peaks, implying the high propensity of these residues to form the allosteric binding pocket. Moreover, these NTD residues are aligned with high binding pocket propensity in each of the three closed protomers.

**Figure 6.**
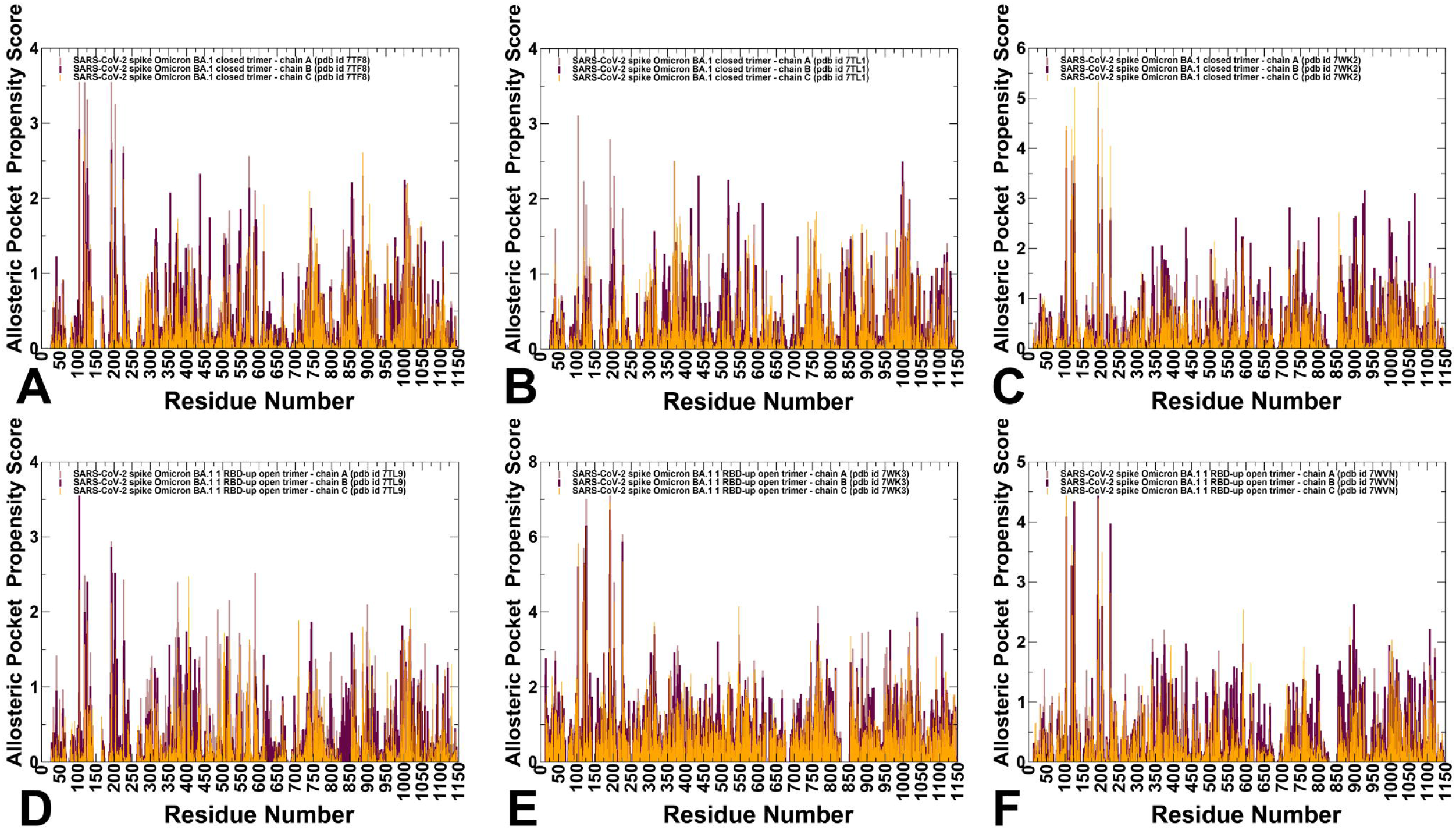
The residue-based pocket propensity distributions for the SARS-CoV-2 S Omicron BA.1 structures. (A-C) The allosteric pocket propensity profile for the closed S-BA.1 trimers (pdb id 7TF8 panel A, pdb id 7TL1 panel B, and pdb id 7WK2 panel C). (D-F) The allosteric pocket propensity profile for the open S-BA.1 trimers (pdb id 7TL9 panel D, pdb id 7WK3 panel E, and pdb id 7WVN/7WVO panel F). The profiles in each structure are shown for the three protomers, protomer A in light brown bars, protomer B in maroon bars, and protomer C in orange bars.

**Figure 7.**
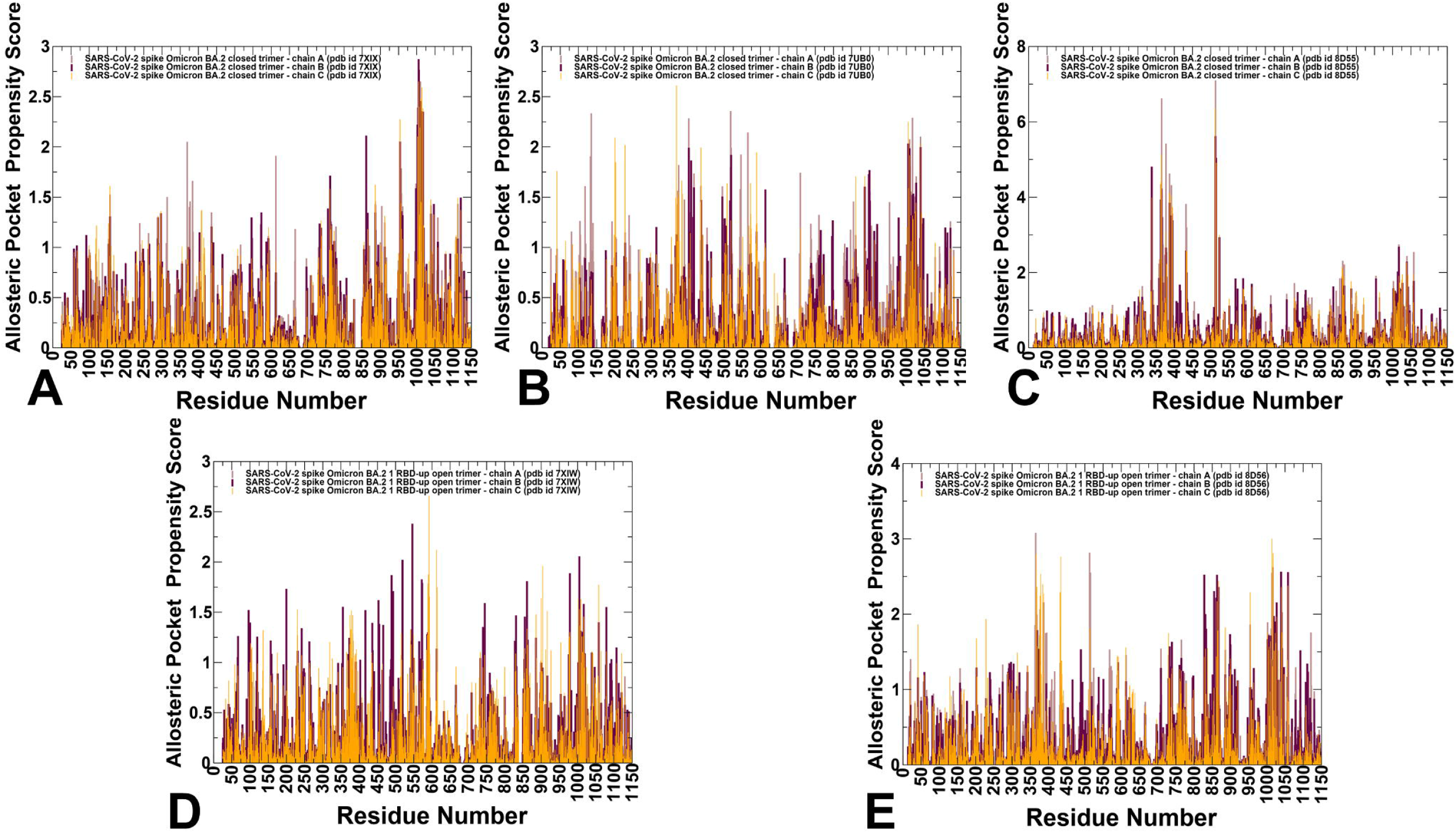
The residue-based pocket propensity distributions for the SARS-CoV-2 S Omicron BA.2 structures. (A-C) The allosteric pocket propensity profile for the closed S-BA.1 trimers (pdb id 7XIX panel A, pdb id 7UB0 panel B, and pdb id 7WK2 panel C). (D-F) The allosteric pocket propensity profile for the open S-BA.1 trimers (pdb id 7TL9 panel D, pdb id 7WK3 panel E, and pdb id 7WVN/7WVO panel E). The profiles in each structure are shown for the three protomers, protomer A in light brown bars, protomer B in maroon bars, and protomer C in orange bars.

The predicted NTD pocket residues overlap with the experimentally known NTD supersite formed by residues 14–20, residues 140–158 (the supersite b-hairpin), and residues 245–264 (the supersite loop). NTD-neutralizing monoclonal antibodies target the same antigenic supersite, providing examples of convergent solutions to NTD-targeted monoclonal antibody neutralization [63,149]. The dominance of these NTD favorable pocket propensity scores is also maintained for the 1RBD-up S-BA.1 conformations (Figure 6D-F). Smaller but notable peaks are associated with clusters of residues in the RBD region, residues from CTD1 and local clusters in the S2 subunit corresponding to HR1 and CH regions (Figure 6A-C). Interestingly, while the distribution of binding pocket propensities for the S-BA.1 forms is mainly determined by the residues from dynamic S1 subunit, a significant fraction of the profile is associated with HR1 and CH regions of the rigid S2 subunit, thereby indicating that an appreciable level of plasticity is present in S2 regions, giving rise to the broader accumulation of dynamic cryptic pockets.

The BA.1 and BA.2 S proteins differ in the NTD with only the G142D substitution shared between the two Omicron variants. The S-BA.2 NTD lacks the H69-V70 deletion (ΔH69-V70) that is present in BA.1, as well as in the Alpha (B.1.1.7). The BA.2 NTD also lacks the deletion of residues 143–145 and the insertion of three residues at position 214. The distribution of network-weighted binding pocket propensities in the closed S-BA.2 conformations showed a broader distribution of favorable pocket sites in the S1-S2 interfaces and inter-protomer regions of S2 subunit (Figure 7A-C). Notably, the high distribution peak corresponding to the NTD regions in the S-BA1 conformations is conspicuously absent for the S-BA.2 ensembles, suggesting that variant-specific remodeling of the NTD may lead to potential alterations and redistribution of cryptic sites. In the S-BA.2 conformations, the NTD residues featured moderate binding pocket propensity values, indicating that the NTD supersite may become partly remodeled and become less favorable for ligand binding. These findings may have particular importance in light of the mechanism in which Omicron and other VOC ‘s harbor frequent mutations within the NTD supersite, pointing to antibody-mediated evolutionary pressure to leverage this domain for antibody evasion [149].

At the same time, the largest peaks in the closed and open S-BA.2 conformations are shifted towards S1-S2 inter-domain regions and particularly residue clusters from HR1 and CH regions of the S2 subunit (Figure 7). These findings signaled that Omicron variants may have evolved to increase conformational plasticity and adaptability of the S2 regions, while maintaining functionally significant changes in the NTD and RBD regions. In other words, our results suggested a broadening of residue pocket propensities in the S-BA.2 conformations, leading to a more heterogeneous allocation of cryptic binding pockets across both S1 and S2 regions.

### 3.5 Deciphering Anatomy of Cryptic Binding Pockets in the S-BA.1 and S-BA.2 Conformational Ensembles

We performed a detailed quantitative analysis of the top ranked pockets in the S-BA.1 and S-BA.2 ensembles to connect computational predictions guided by allosteric role of the binding residues with the existing experimental data and functional knowledge of S protein mechanisms. To understand the predictive ability of the approach and decipher functional role of the cryptic pockets, we started by comparing complete structural maps of all identified pockets versus the top ranked cryptic sites for the ensembles of different S-BA.1 structures (Figure 8) and S-BA.2 structures (Figure 9). First, the structural projections of all pockets showed that cryptic cavities could be found in both S1 and S2 regions. Moreover, the identified binding pockets form a dense network of overlapping sites which may have some relevance for propagating signals in the S protein. However, the “universe” of the computed S1 binding pockets does not necessarily mean that all of them or even the large fraction of them have functional significance and could represent targetable structural elements for antibody and vaccine design purposes. Based on this premise, we then compared the top ranked cryptic sites for different S-BA.1 (Figure 8) and S-BA.2 conformations (Figure 9). The top 5-7 ranked cryptic sites in the S-BA.1 are generally consistent between different BA.1 structural ensembles (Figure 8A-E). Indeed, for all S-BA.1 conformations, the top ranked NTD binding pocket is found for each of the protomer. We also identified several other highly ranked and functionally significant cryptic sites formed at the inter-domain and inter-protomer interfaces. It could be seen that the top ranked pockets in the S-BA.1 conformations favors the S1 and S1-S2 regions, with only a fraction of the binding sites in the dense S2 regions (Figure 8A-E).

**Figure 8.**
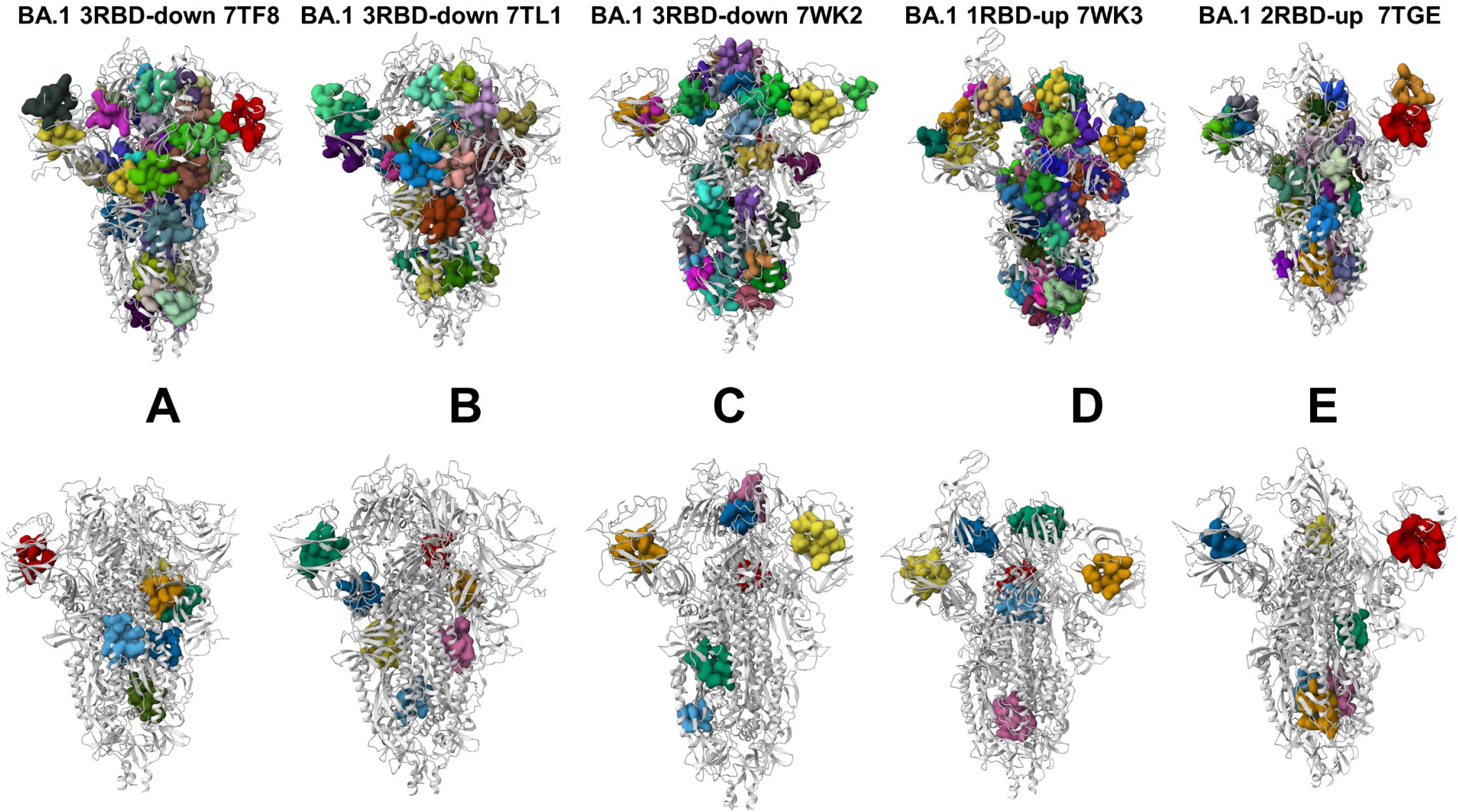
Structural maps of all identified pockets versus the top ranked cryptic sites for the ensemble of different S-BA.1 structures. Structural map of the cryptic pockets for the closed S-BA.1 trimer, pdb id 7TF8 (A), closed BA.1 trimer, pdb id 7TL1 (B), closed S-BA.1 trimer, pdb id 7WK2 (C), open BA.1 trimer, pdb id 7WK3 (D) and open BA.1 trimer, pdb id 7TGE.

**Figure 9.**
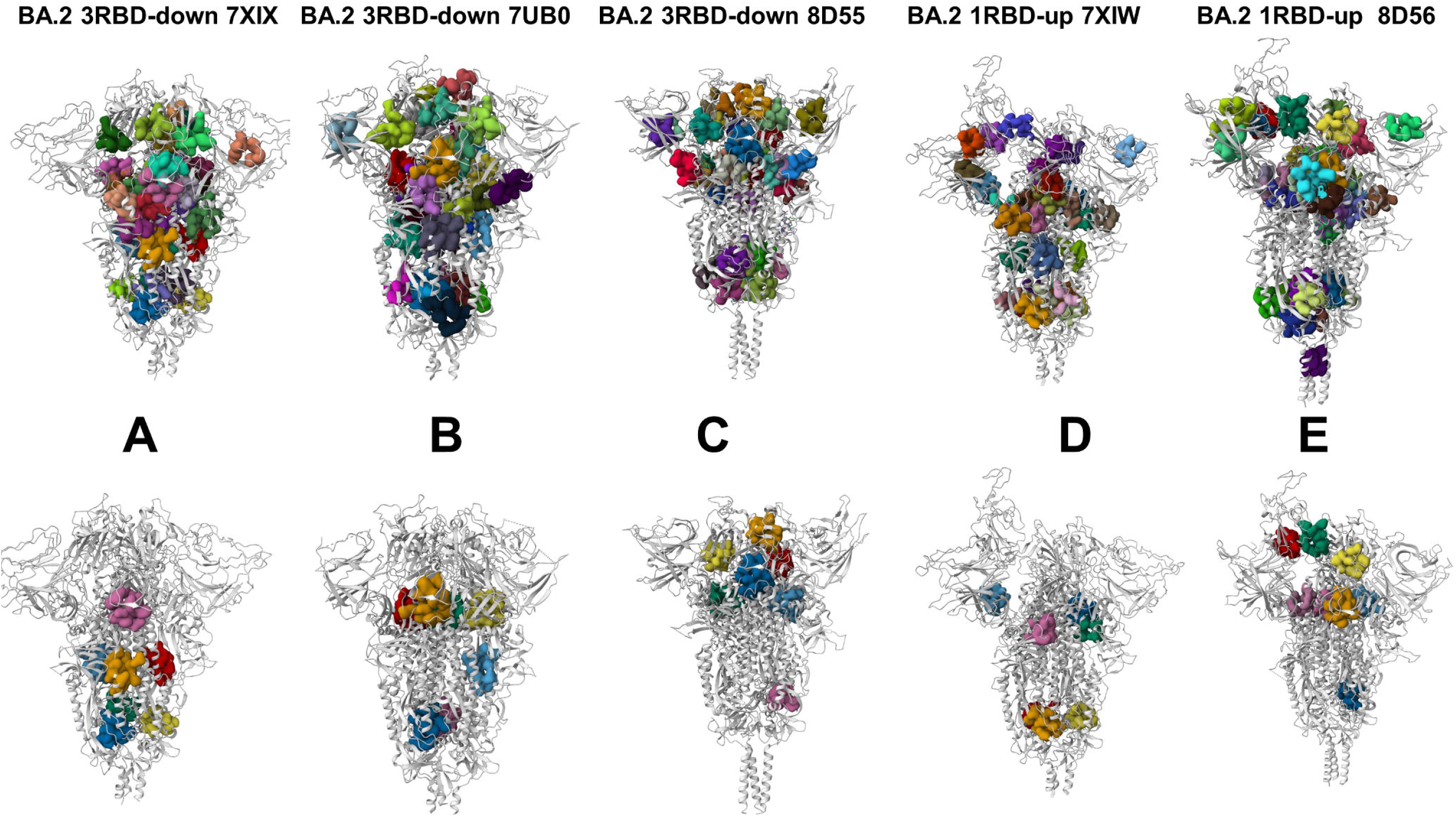
Structural maps of all identified pockets versus the top ranked cryptic sites for the ensemble of different S-BA.2 structures. Structural map of the cryptic pockets for the closed S-BA.2 trimer, pdb id 7XIX (A), closed BA.2 trimer, pdb id 7UB0 (B), closed S-BA.2 trimer, pdb id 8D55 (C), open BA.2 trimer, pdb id 7XIW (D) and open BA.2 trimer, pdb id 8D56.

A similar heterogenous picture of the all predicted pockets (Figure 9A-E). However, the allosteric-guided ranking of the top cryptic pockets showed interesting differences between BA.1 and BA.2 ensembles. We noticed that the NTD regions in the S-BA.2 ensembles did not harbor the most probable cryptic pockets, while these pockets were found in the RBD, S1-S2 and S2 regions (Figure 9A-E). The distributions of the top pockets varies between different S-BA.2 closed conformations. Interestingly, in all BA.2 structures the top ranked pockets included RBD and RBD-RBD regions. Although the distribution of the top ranked binding sites reflected heterogeneity of both BA.1 and BA.2 trimers, BA.1 pockets emerge more consistently in the different BA.1 structures. Given the observed structural rigidity of the S-BA.1 closed trimers, the emergence of highly ranked cryptic sites in the S2 subunit pointed to some underappreciated plasticity of the S2 regions (Figure 8). We also noticed that the ensemble of predicted pockets in the S-BA.1 trimers is more interconnected pointing to a possibility of rapid communication between the sites. A more dynamic character of the S-BA.2 open trimers produced several islands of disconnected pocket clusters as many other dynamically formed cavities do not persist in the course of simulations.

To validate the predictions and rationalize possible functional role of the top ranked allosteric sites, we followed by presenting a detailed pocket analysis of the S-BA.1 closed trimer conformations and the top six cryptic pockets determined from consensus in this analysis (Figure 10). Strikingly, the top predicted NTD binding pocket (Figure 10, pocket 1) is precisely aligned with the experimentally discovered binding site that is lined by hydrophobic residues I101, W104, I119, V126, M177, F192, F194, I203, and L226 which form the interactions with the ligand in the cryo-EM structure [61,62]. In fact, experimental studies showed that binding of polysorbate 80 (PS80) and heme metabolites biliverdin and bilirubin to this cryptic NTD site can alter the epitope and modulate the antibody response [60–62,112]. The experimentally observed immune evasion through recruitment of small heme metabolite molecules [60–62] revealed the functional significance of the top predicted pocket (Figure 10, pocket 1). Importantly, one effective NTD-targeting antibody, P008_056 competes with small molecule biliverdin for binding to the NTD pocket allosterically by inducing conformational changes in the loop 175-185 [61], while another effective NTD antibody PVI.V6-14 [62] binds directly to the NTD hydrophobic pocket identified in our analysis. The network-based screening of the binding pockets not only captured this experimentally validated cryptic pocket with all functional residues present but also ranked it as the most probable for ligand binding in the closed and open forms of the S-BA.1 trimer (Figure 10). The latest studies highlighted conformational plasticity of the NTD regions where mutations and/or deletions not only change the architecture, but also alter the surface properties, leading to remodeling of the binding pockets and major antigenic changes in the NTD supersite and loss of antibody binding [63].

**Figure 10.**
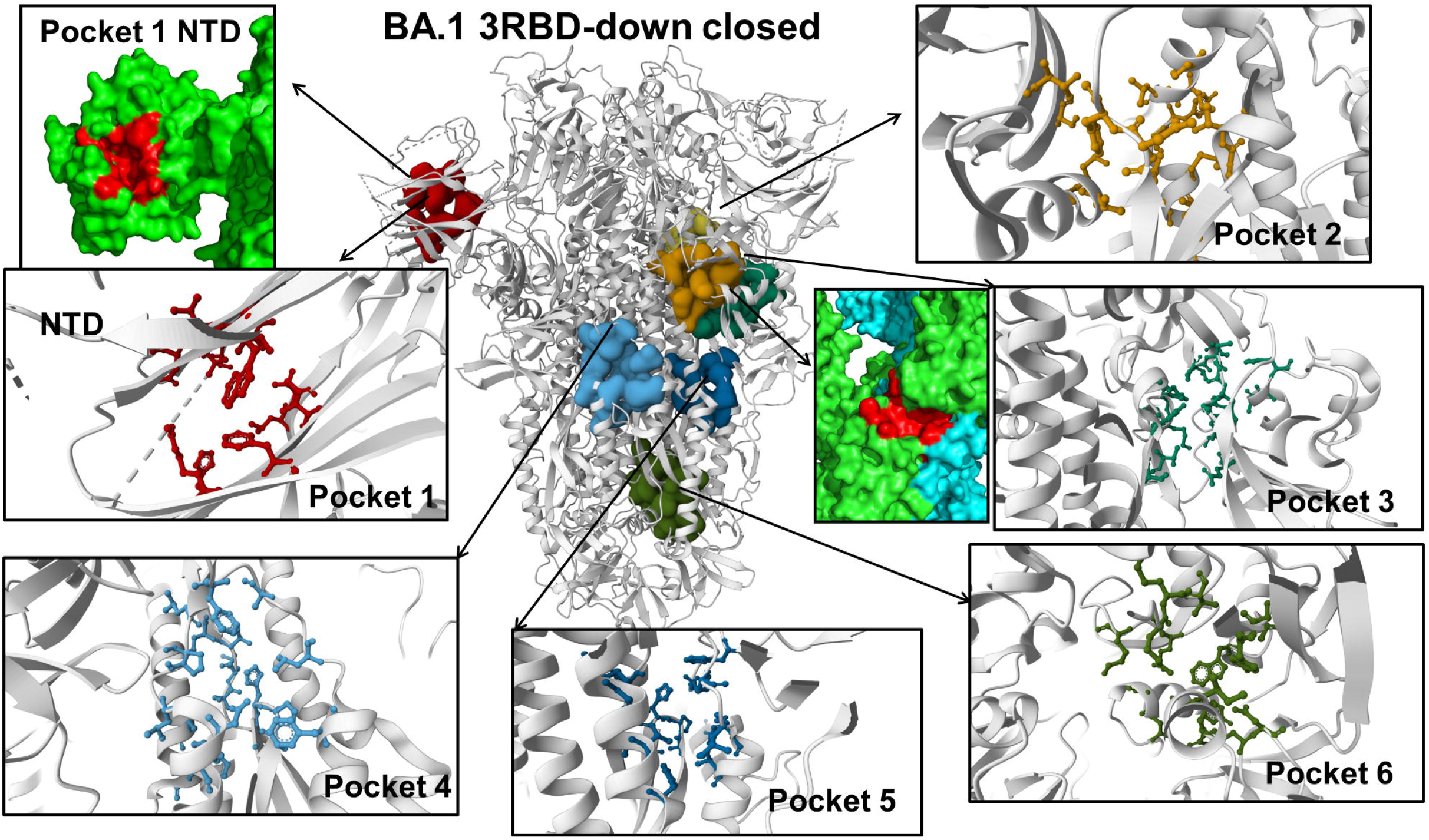
Structural map and residue-based close-ups of the top ranked cryptic sites for the ensemble of the S-BA.1 closed trimers. The BA.1 closed trimer (pdb id 7TF8) is shown in ribbons and the top ranked allosteric sites are in surface. Pocket 1 (residues 101, 104, 119, 121, 126, 128, 170, 172, 190, 192, 194, 203, 205, 207, 226, 227 in each of the protomer). Pocket 1 is also highlighted in red surface with NTD in green. Pocket 2 (residues B_274, B_302, B_304, B_316, B_317, B_318, B_50, C_738, C_739, C_750, C_753, C_754, C_756, C_757, C_760, C_761, C_764). Pocket 3 (residues B_551, B_588, B_589, B_590, B_591, B_592, B_593, B_594, B_613, B_622, B_623, B_625, C_735, C_737, C_740, C_837, C_841, C_854, C_855, C_857, C_859). Pocket 3 is also shown in red surface at the inter-protomer interface of two protomers (in green and cyan surface). Pocket 4 (residues B_1056, B_1057, B_1058, B_1059, B_730, B_731, B_732, B_733, B_778, B_782, B_823, B_828, B_831, B_833, B_861, B_863, B_865, B_867, B_870). Pocket 5 (residues C_1056, C_1057, C_1058, C_1059, C_729, C_730, C_731, C_774, C_778, C_782, C_823, C_828, C_833, C_863, C_865, C_867, C_870). Pocket 6 (residues B_1037, B_1039, B_1040, B_1046, B_1047, B_1107, B_1108, B_909, B_910, C_1036, C_1038, C_886, C_890, C_904, C_907, C_908)

Another highly probable pocket is formed at the inter-protomer interface by residues T274, T302, S316, F318 of one protomer and residues D737, C738, T739, S750, L753 L754, Y756, C760, T761 and K764 of the neighboring protomer (Figure 10, pocket 2). Importantly, this pocket is anchored by the key hinge position F318 on one protomer and N764 Omicron mutational site, being stabilized through the interprotomer bridges N317-D737, R319-D737, and R319-T739. Our pocket discovery analysis highlighted as one of the five top binding pockets a cryptic site at the inter-protomer interface formed by residues V551, T588, P589, C590, S591, F592, G593, Q613, G614 of one protomer and residues D737, M740, Y837, L841, K854, F855, K856, G857 and T859 of the neighboring protomer (Figure 10, pocket 3).

Strikingly, this cryptic site provides a fairly small and deep pocket formed by critically important inter-protomer residues. Indeed, positions S591 and F592 are conserved structurally stable hinge sites of collective motions that can modulate both the inter-protomer and inter-domain changes. This binding pocket is located immediately next to the ordered FPPR motif (residues 823-858) that engages in allosteric cross-talk with the RBD regions [150]. Moreover, residues K854, F855 and K856 are critical inter-protomer sites involved in the inter-protomer contacts (such as N856K-D568, N856K-T572) and changes in these positions can modulate the shifts between the open and closed sites. For example, the quadruple mutant (A570L/T572I/F855Y/N856I) introducing modifications in these positions can shift the equilibrium away from the closed-down state [151]. The emergence of this cryptic pocket and its high ranking in our analysis is also due to both the network-mediating allosteric properties of the respective residues and their high ligand binding propensities.

In contrast to the large accumulation of mutations within the RBD, the S2 fusion subunit has remained highly conserved among variants but Omicron mutations targeted this subunit as well. The recently developed broad-spectrum fusion inhibitors and candidate vaccines target the conserved elements in the S2 subunit, including the fusion peptide, stem helix, and HR1 and HR2 regions [112]. Our analysis identified a cryptic binding pocket in the S2 subunit formed by S1037, R1039, V1040, G1046, Y1047, R1107, N1108, I909, G910 of one protomer and residues W886, A890, Y904, G908 of the adjacent protomer (Figure 10, pocket 6). This pocket is formed deeply in the S2 subunit and formed by residues of the C-terminal β-hairpin (β_49_–β_50_) (residues 1045-1076) located downstream of the CH region. Interestingly, in several closed S-BA.1 structures (pdb id 7TL1, 7WK2) (Figure 8B,C) and one S-BA.1 open conformation (pdb id 7WK3) (Figure 8D) we also identified as one of the top pickets the free fatty acid (FA) binding site [152] that bridges two RBDs of adjacent protomers (residues 368, 369, 372, 374, 377, 379, 384, 387, 436, 437, 440, 503, 506 of one RBD protomer and residues 493,405, 501, 502, 503, 504, 505 of another RBD protomer) (Figure 8C). In agreement with structural studies [152,153], we found that the RBD allosteric pocket may partly collapse in the open BA.1 conformations. However, this pocket was still observed in one of the open BA.1 conformations. These observations indicated that the distribution of cryptic pockets may be sensitive to the conformational landscapes of S trimers. Indeed, even for structurally similar and stable BA.1 closed trimers, the conserved allosteric pocket between two RBDs may partly vary among different structures (Figure 8).

The pocket analysis of the S-BA.2 conformations revealed a different picture and showed evidence of variant-specific distribution of putative allosteric sites (Figures 9,11). First, the NTD binding pocket was no longer ranked as the top allosteric site as conformational plasticity of the NTD regions and supersite is increased in the S-BA.2 conformations causing appreciable remodeling of the NTD binding pocket (Supplementary Materials, Figure S1). Indeed, a comparison of the NTD binding sites in different BA.1 and BA.2 conformations revealed NTD rearrangements manifesting in fragmentation and partial occlusion of the predicted pocket in the closed and open BA.2 ensembles (Supplementary Materials, Figure S1).

**Figure 11.**
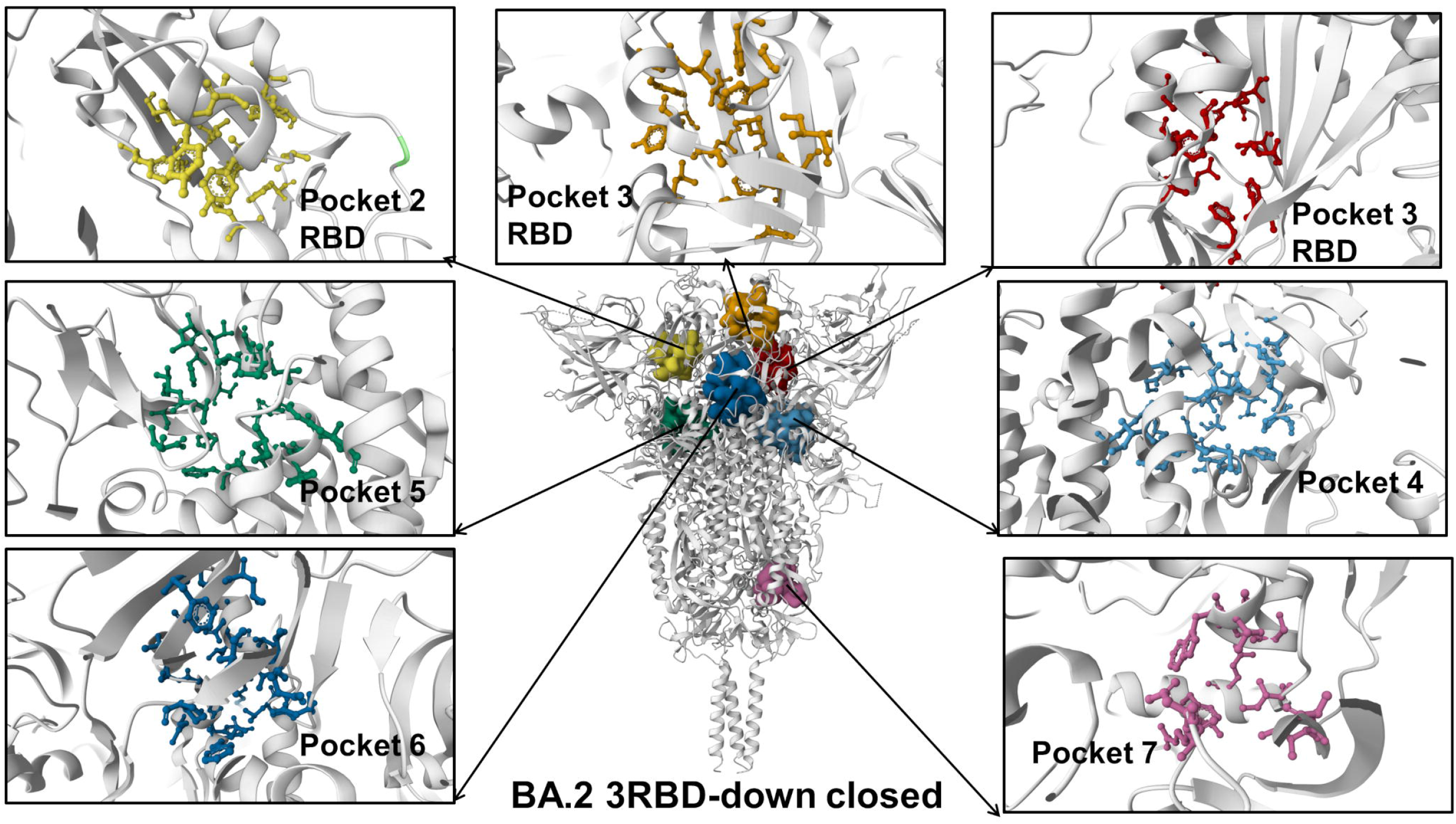
Structural map and residue-based close-ups of the top ranked cryptic sites for the ensemble of the S-BA.2 closed trimers. The BA.2 closed trimer (pdb id 8D55) is shown in ribbons and the top ranked allosteric sites are in surface. Pocket 1 (residues A_338, A_363, A_365, A_368, A_374, A_377, A_387, A_392, A_395, A_432, A_434, A_513, A_515, protomer A). Pocket 2 (residues C_338, C_358, C_363, C_365, C_368, C_369, C_377, C_387, C_392, C_395, C_432, C_434, C_513, C_515, protomer C). Pocket 3 (residues B_338, B_342, B_358, B_363, B_365, B_368, B_369, B_377, B_387, B_392, B_395, B_432, B_434, B_513, B_515, protomer B). Pocket 4 (residues A_541, A_546, A_548, A_549, A_568, A_570, A_572, A_573, A_574, A_587, A_588, A_589, C_1000, C_740, C_741, C_742, C_744, C_745, C_845, C_852, C_855, C_856, C_966, C_975, C_976, C_977, C_978). Pocket 5 (residues B_1000, B_740, B_741, B_742, B_744, B_745, B_855, B_856, B_966, B_975, B_976, B_977, B_978, C_541, C_546, C_547, C_548, C_549, C_568, C_572, C_573, C_587, C_589). Pocket 6 (residues A_1000, A_740, A_741, A_742, A_744, A_745, A_855, A_856, A_966, A_975, A_976, A_977, A_978, B_541, B_546, B_547, B_548, B_549, B_568, B_570, B_572, B_573, B_574, B_587, B_589). Pocket 7 (residues A_1107, A_1108, A_712, A_713, C_884, C_885, C_886, C_887, C_896, C_901, C_904).

Our findings are consistent with recent studies of BA.1 and BA.2 variants showing that the neutralizing activity of anti-NTD monoclonal antibodies was markedly impaired due to the BA,2 mutation of G142D and del 143-145 in the NTD [40,52]. The NTD of BA1 is more remodeled compared to BA.2, presenting three substitutions A67V, T95I, and G142D, 5 deleted residues (Δ69–70 and Δ143–145), and three inserted residues. As a result, NTD binding antibodies isolated from BA.2 do not neutralize BA.1 [154]. According to our observations, these mutational differences in the NTD between BA.1 and BA.2 can affect conformational landscape of the NTD regions leading to more adaptable and stable NTD binding pocket in which respective contributing residues have significant allosteric potential. This NTD binding pocket is remodeled and is less stable in the S-BA.2 conformations (Figure 9).

The sensitivity of the cryptic pockets formation to the conformational dynamics is far more pronounced for the S-BA.2 structures (Figure 9). While pocket analysis in some of the BA.2 conformations did not detect the RBD LA pocket (Figure 9A,B), we found that this experimentally known cryptic site emerged as the most probable allosteric site for the ensemble of the closed BA.2 trimer (pdb id 8D55, Figure 9C) and was also found in the open BA.2 conformation (pdb id 8D56, Figure 9E). The top ranked LA pocket in the BA.2 states is formed by RBD residues F338, I358, A363, Y365, L368, Y369, F377, L387, F392, V395, C432, I434, L513, F515 on each protomer (Figure 11, pockets 1-3 for protomers A, C and B).

A mapping of the LA pocket RBD residues displaying high ligand binding propensities and mediating potential demonstrated that these sites tend to occupy other well-defined pockets on the RBD surface in the S-BA.2 conformations that form “pocket-based” pathways connecting the allosteric site with the RBD binding interface patches (Supplementary Materials, Figure S2). The greater conformational sensitivity of the predicted cryptic pockets in the S-BA.2 structures is consistent with the dynamics analysis showing that the BA.2 forms are more flexible. In addition, these important findings corroborate with another computational study [155] showing that the RBD–RBD interface is less tightly packed in the closed and open BA.2 trimers. Our results confirm the notion that efficiency of allosteric LA binding to the RBD site can be determined by the Omicron variants [153,155,156]. In particular, we found that the LA allosteric pocket can be transiently formed in some open BA.2 conformations (Figure 9E) but not detected in other BA.2 open states (Figure 9D). Our findings that some BA.2 closed conformations (such as locked closed trimer from pdb id 8D55, Figure 9C) could prominently feature the LA binding pocket as the top ranked site can be better understood in the context of recent functional studies [156] showing that compact locked 3 RBD-down S trimer is stabilized by LA occupying a bipartite binding site composed of two adjacent RBDs in the trimer. According to this elegant structural study, LA binding to the RBD can induce compaction of the three RBDs, sharing characteristic features with the locked S trimer [156]. Indeed, our predictions of the cryptic pockets in the ensemble of the locked S-BA.2 trimer conformation (Figure 9C) yielded the LA pocket as the most dominant.

Common with S-BA.1 trimers, one of the highly ranked pockets in the closed BA.2 is formed at the inter-protomer interface by residues F541, L546, D568, A570, T572, T588 and P589 of one protomer and residues D737, M740, Y837, L841, K854, F855, K856, G857 and T859 of the neighboring protomer (Figure 11). A close-up comparison of this inter-protomer site between closed S-BA.1, closed S-BA.2 and open S-BA.2 conformations showed a well-defined and conserved binding pocket in the closed states (Supplementary Materials, Figure S3). This pocket has a significant functional importance as it is formed at the inter-protomer juncture anchored by known allosteric hotspot positions A570, T572, K854, F855, K856 in which mutations can alter conformational equilibrium. As a result, it is likely that binding of a small molecule to this pocket may affect the large conformational changes and effectively prevent S protein from adopting an open state.

The distributions for the S-BA.2 closed trimers showed marked preference towards S2 subunit pockets, particularly CH region residues as having the largest propensities. One of the most probable pockets is formed by S1037, R1039, V1040, G1046, Y1047, R1107, N1108, I909, G910 of one protomer and residues W886, A890, Y904, G908 of the adjacent protomer (Figure 11). This pocket was also ranked highly in the S-BA.1 conformations (Figure 10, pocket 6). Another functionally relevant cryptic pocket formed by S2 residues of all three protomers (residues 1024, 1027, 1039, 1042 on each protomer) presents a target for Umifenovir (Arbidol)—a broad-spectrum anti-influenza drug—that binds to the hydrophobic cavity of the trimer and inhibits post-fusion conformational transition related to membrane fusion [157,158]. It was proposed that Arbidol interacts with residues in the central CH-HR1 helix residues K776, E780, K947, E1017, R1019, S1021, N1023, L1024 and T1027 of the S2 trimerization domain [157,158] and can interfere with the trimerization of the S protein, which is critical for virus entry. We also found another highly ranked pocket composed of residues on adjacent protomers and overlapping with the highly conserved, conformational hinge epitope spanning residues 980–1006 of the S protein, at the apex of the S2 domain [159]. Our analysis uncovered the presence of this hidden pocket in the closed form, but the hinge epitope is only accessible to antibody binding in the open form. The discovery of these pockets that overlap or immediately adjacent to the S2 hinge epitope [159] are important in light of findings that antibody evasion may occur through dynamics changes in the neighboring regions that impact hinge epitope accessibility. The hinge epitope and neighboring cryptic pockets are protected by tightly packing the RBDs in the down state, but access to these pockets may be modulated through global dynamic changes and long-range couplings between S1 and S2 subunits can affect formation and stability of the cryptic binding sites.

## 4. Discussion

Our study leveraged the growing wealth of closed and open structures for the S-BA.1 and S -BA.2 trimers to explore conformational landscapes of these S proteins and examine the evolution and character of cryptic binding pockets in these ensembles. Despite considerable structural similarities between the closed 3RBD-down structures, the conformational landscapes revealed some specific characteristics that are translated into rich distribution of the identified cryptic pockets. Using allosteric-based network ranking of the binding pockets, we recovered all experimentally known allosteric sites and discovered significant variant-specific differences in the distribution of cryptic binding sites in the BA.1 and BA.2 trimers. While both closed and open S-BA.1 trimers featured as highly ranked NTD supersite region [63], the conformational dynamics of the S-BA.2 trimers could partly mask the NTD cryptic region and present previously underappreciated cryptic binding pockets in the inter-protomer interface and hinge regions of the S2 subunit. The in-depth analysis of the conformation-specific evolution of the NTD pockets highlighted the notion that mutational changes in the NTD between BA.1 and BA.2 are translated into conformational adaptability and variant-specific remodeling of the NTD site. Mutational changes in the BA.1 variant can induce druggable NTD pocket that is preserved and ranked as the most probable in both closed and open BA.1 conformations. This result is particularly interesting suggesting that despite conformational plasticity that is afforded in the NTD, this allosteric pocket is preserved and accessible for ligand binding in the BA.1 conformations. At the same time, our findings suggested that distinct mutational signature in the NTD of BA.2 trimers may result in considerable remodeling and fragmentation of the NTD binding pocket. These revelations provide support to the recent data suggesting that the NTD can serve as an adaptable antigenic surface capable of unlocking and redistributing cryptic binding pockets which may enable the efficient immune escape away from the RBD regions [50]. Interestingly, our studies highlighted that the NTD allosteric binding sites harbor variable regions and can undergo considerable remodeling between Omicron variants [61] whereas evolutionary conservation of the LA-binding pocket is the Omicron variants [156] can co-exist with partial restructuring and adaptability of the RBD-RBD. Our predictions of conformation and variant-specific preferences of the LA pocket are in line with studies of LA binding with the S protein showing scenarios of dynamic opening-closing of the pocket through the gating helix motions or opening of the pocket while interacting with LA [156]. We argue that the LA pocket opening/closing is linked with the conformational plasticity of the S trimers, particularly for the BA.2 variant. As a result, LA binding may occur through a conformational selection of a closed or open conformational state in which the RBD-RBD pocket is accessible followed by LA-induced stabilization of the locked closed form. The observed variant-sensitive adaptability of the RBD-RBD allosteric site could also affect how structural changes in the RBD are propagated to other functional sites, while preserving the general topology of the LA binding pocket in spike.

Understanding of the interplay of conformational dynamics changes induced by Omicron variants and identification of cryptic dynamic binding pockets in the S protein are of paramount importance in COVID-19 research as exploring broad-spectrum antiviral agents to combat the emerging variants is imperative. In contrast to significant accumulation of mutations within the RBD, the S2 fusion subunit has remained highly conserved among variants with only several mutational sites targeted by Omicron variants. Identifying druggable sites in the S2 subunit can present a new and previously underappreciated opportunity for therapeutic intervention given the increasing number of fusion inhibitors and candidate vaccines that target the conserved elements in the S2 subunit [112,113]. The results of our study suggested that despite general rigidity of the S2 regions in comparison with more dynamic S1 subunit, there is still an appreciable level of conformational adaptability in the S2, resulting in significant number of dynamic cryptic pockets in the S2. In particular, our data pointed to several conserved pockets in the HR1 and CH regions of the rigid S2 subunit, thereby indicating that an appreciable level of plasticity is present in S2 regions, giving rise to the broader accumulation of dynamic cryptic pockets. The results of this study provided molecular rationale and support to the experimental evidence that the acquisition of functionally balanced substitutions that optimize multiple fitness tradeoffs between immune evasion, host binding affinity and sufficient conformational adaptability might be a common strategy of the virus evolution and serve as a primary driving force behind the emergence of new Omicron subvariants.

## 5. Conclusions

In the current study we explore conformational landscapes and characterize the universe of cryptic binding pockets in multiple open and closed functional spike states of the Omicron BA.1 and BA.2 variants. The conformational dynamics and network analysis confirmed that S-BA.1 and S-BA.2 trimers may display different conformational plasticity and dynamic changes, allowing for modulation for opening and closing cryptic binding pockets in the S1 and S2 subunits. By using a combination of atomistic simulations, dynamics network analysis, and allostery-guided network screening of cryptic pockets in the conformational ensembles of BA.1 and BA.2 spike conformations, we demonstrated that the proposed approach would recover the experimentally known allosteric sites in the NTD, RBD regions as well as targetable S2 regions as highly ranked top binding pockets. The results of this study indicate that while both closed and open S-BA.1 trimers featured as highly ranked NTD supersite region, the conformational dynamics of the S-BA.2 trimers could mask the NTD cryptic region and present previously underappreciated cryptic binding pockets in the inter-protomer interface and hinge regions of the S2 subunit. We found that mutational differences in the NTD between BA.1 and BA.2 can affect conformational landscape of the NTD regions leading to more adaptable and stable NTD binding pocket in the BA.1 structures, while the NTD binding pocket is remodeled and is less stable in the S-BA.2 conformations. The central result of this analysis is discovery of variant-specific differences in the distribution of cryptic binding sites in the BA.1 and BA.2 trimers, suggesting that small variations could lead to different preferences in allocation of druggable sites. The predicted conformation and variant-specific preferences of the RBD allosteric pocket agree with the observed dependency of LA binding to the RBD site between the Omicron variants and binding preference towards locked closed S conformations. The determined cryptic binding pockets at the inter-protomer regions and in the functional regions of the S2 subunit such as HR1-HR2 bundle and stem helix region, are consistent with the role of the pocket residues in modulating conformational transitions and antibody recognition. Of particular interest is the detection of highly probable pockets in the S2 regions known to be allosterically linked with the RBD movements. The results of this study are particularly significant for understanding the universe of cryptic bindings sites detection of conformation-dependent variant-specific preferences for conserved druggable pockets. Targeted ligand screening in the predicted druggable sites can allow for design of variant-specific modulators of the S activity and ligand-induced stabilization of specific conformational states, thus providing chemical tools for systematic probing of the S functions and binding. The results of this study provided better understanding of the interplay between conformational plasticity and evolution of novel druggable cryptic pockets in the S trimers, suggesting diversity of targetable regions and variant-specific preferences of allosteric pockets. This may enable engineering of efficient and selective modulators that could rationally target a complex functional landscape of virus transmissibility.

## Supplementary Materials

The following supporting information can be downloaded at: www.mdpi.com/xxx/s1, Figure S1: A comparison of the predicted NTD binding pockets in the closed BA.1 trimer structures, BA.2 closed trimer structures and BA.2 open trimer conformations; Figure S2: A comparison of the predicted RBD binding pocket for the BA.1 closed conformation and BA.2 closed conformations; Figure S3: A comparison of the predicted inter-protomer cryptic pocket in the BA.1 closed trimers, BA.2 closed trimers and BA.2 open trimer conformations.

## Author Contributions

Conceptualization, G.V.; methodology, G.V.; software, G.V., M.A. and G.G; validation, G.V.; formal analysis, G.V., M.A. and G.G.; investigation, G.V.; resources, G.V., M.A. and G.G.; data curation, G.V.; writing—original draft preparation, G.V.; writing— review and editing, G.V., M.A. and G.G.; visualization, G.V.; supervision, G.V.; project administration, G.V.; funding acquisition, G.V. All authors have read and agreed to the published version of the manuscript.

## Acknowledgments

The authors acknowledge support from Schmid College of Science and Technology at Chapman University for providing computing resources at the Keck Center for Science and Engineering.

