## Supplemental Figures S1,S2, S3 for "Ensemble-Based Modeling of the SARS-CoV-2 Omicron BA.1 and BA.2 Spike Trimers and Systematic Characterization of Cryptic Binding Pockets in Distinct Functional States : Emergence of Conformation-Sensitive and Variant-Specific Allosteric Binding Sites"

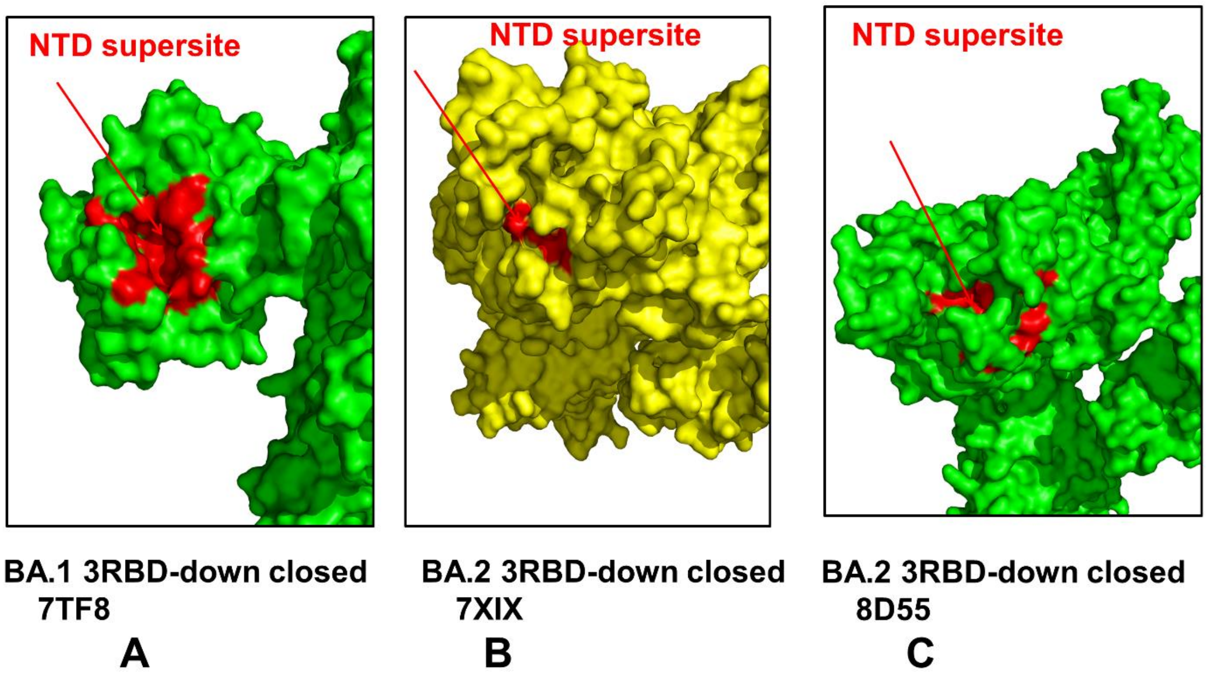


**Figure S1.** A comparison of the predicted NTD binding pockets in the closed BA.1 trimer structures (A), BA.2 closed trimer structures (B) and BA.2 open trimer conformations (C). The S protein is shown in green surface. The predicted NTD binding pocket is shown in red surface and indicated by arrow. The predicted NTD pocket residues overlap with the experimentally known NTD supersite formed by residues 14–20, residues 140–158 (the supersite b-hairpin), and residues 245–264 (the supersite loop).

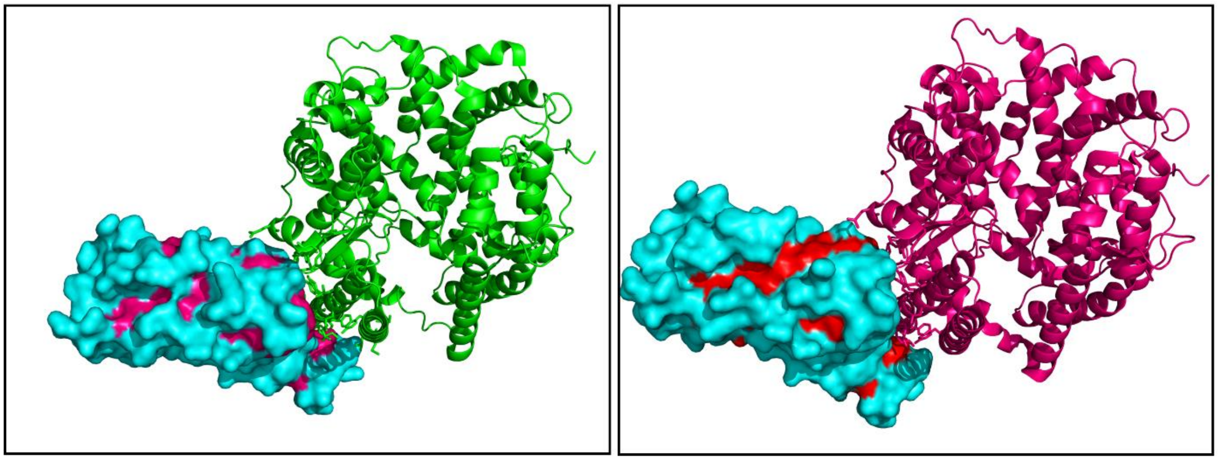

**BA.1 3RBD-down closed**
**7TF8**

**A**

**BA.2 3RBD-down closed**
**7XIX**

**B**

**Figure S2.** A comparison of the predicted RBD binding pocket for the BA.1 closed conformations (A) and BA.2 closed conformations (B). The RBD is shown in cyan surface, the RBD pocket regions formed are shown in pink surface (A) and red surface (B). The host receptor ACE2 is shown in green ribbons (A) and dark pink (B)

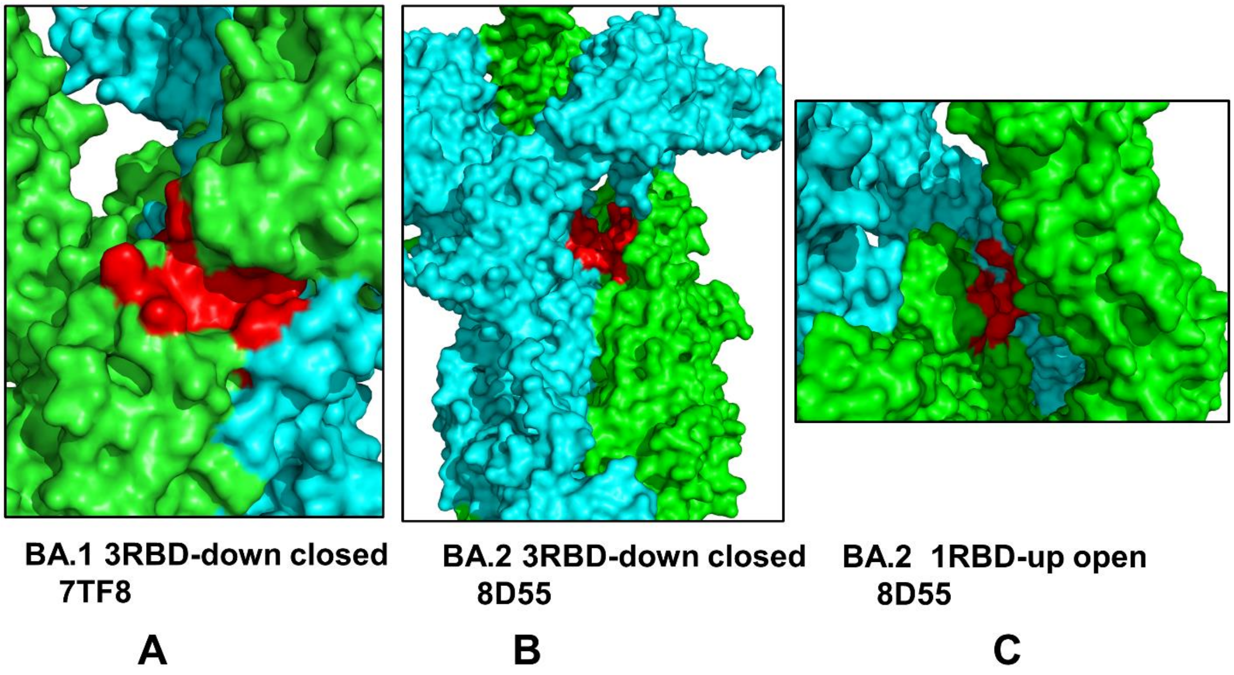


**Figure S3.** A comparison of the predicted inter-protomer cryptic pocket in the BA.1 closed trimers (A), BA.2 closed trimers (B) and BA.2 open trimer conformations (C). The S protein protomers are shown in green and cyan surface. The predicted cryptic pocket is shown in red surface. This cryptic site at the inter-protomer interface is formed by residues V551, T588, P589, C590, S591, F592, G593, Q613, G614 of one protomer and residues D737, M740, Y837, L841, K854, F855, K856, G857 and T859 of the neighboring protomer.
